# Structural basis of odorant recognition by a human odorant receptor

**DOI:** 10.1101/2022.12.20.520951

**Authors:** Christian B. Billesbølle, Claire A. de March, Wijnand J. C. van der Velden, Ning Ma, Jeevan Tewari, Claudia Llinas del Torrent, Linus Li, Bryan Faust, Nagarajan Vaidehi, Hiroaki Matsunami, Aashish Manglik

## Abstract

Our sense of smell enables us to navigate a vast space of chemically diverse odor molecules. This task is accomplished by the combinatorial activation of approximately 400 olfactory G protein-coupled receptors (GPCRs) encoded in the human genome^1–3^. How odorants are recognized by olfactory receptors (ORs) remains mysterious. Here we provide mechanistic insight into how an odorant binds a human olfactory receptor. Using cryogenic electron microscopy (cryo-EM), we determined the structure of active human OR51E2 bound to the fatty acid propionate. Propionate is bound within an occluded pocket in OR51E2 and makes specific contacts critical to receptor activation. Mutation of the odorant binding pocket in OR51E2 alters the recognition spectrum for fatty acids of varying chain length, suggesting that odorant selectivity is controlled by tight packing interactions between an odorant and an olfactory receptor. Molecular dynamics simulations demonstrate propionate-induced conformational changes in extracellular loop 3 to activate OR51E2. Together, our studies provide a high-resolution view of chemical recognition of an odorant by a vertebrate OR, providing insight into how this large family of GPCRs enables our olfactory sense.

## INTRODUCTION

Our sense of smell relies on our ability to detect and discriminate a vast array of volatile odor molecules. The immense chemical diversity of potential odorants, however, poses a central challenge for the olfactory system of all animals. In vertebrates, odorants are detected by olfactory receptors (ORs), which are G protein-coupled receptors (GPCRs) expressed in olfactory sensory neurons (OSNs) projecting from the olfactory epithelium to the olfactory bulb in the brain^1,3^. To detect and discriminate the vast diversity of potential odorants^4^, the OR gene family has expanded dramatically in vertebrate genomes, with some species encoding thousands of OR genes^5^. In humans, the approximately 400 functional ORs constitute half of the broader class A GPCR family (**Fig. 1a**)^6,7^.

**Figure 1.**
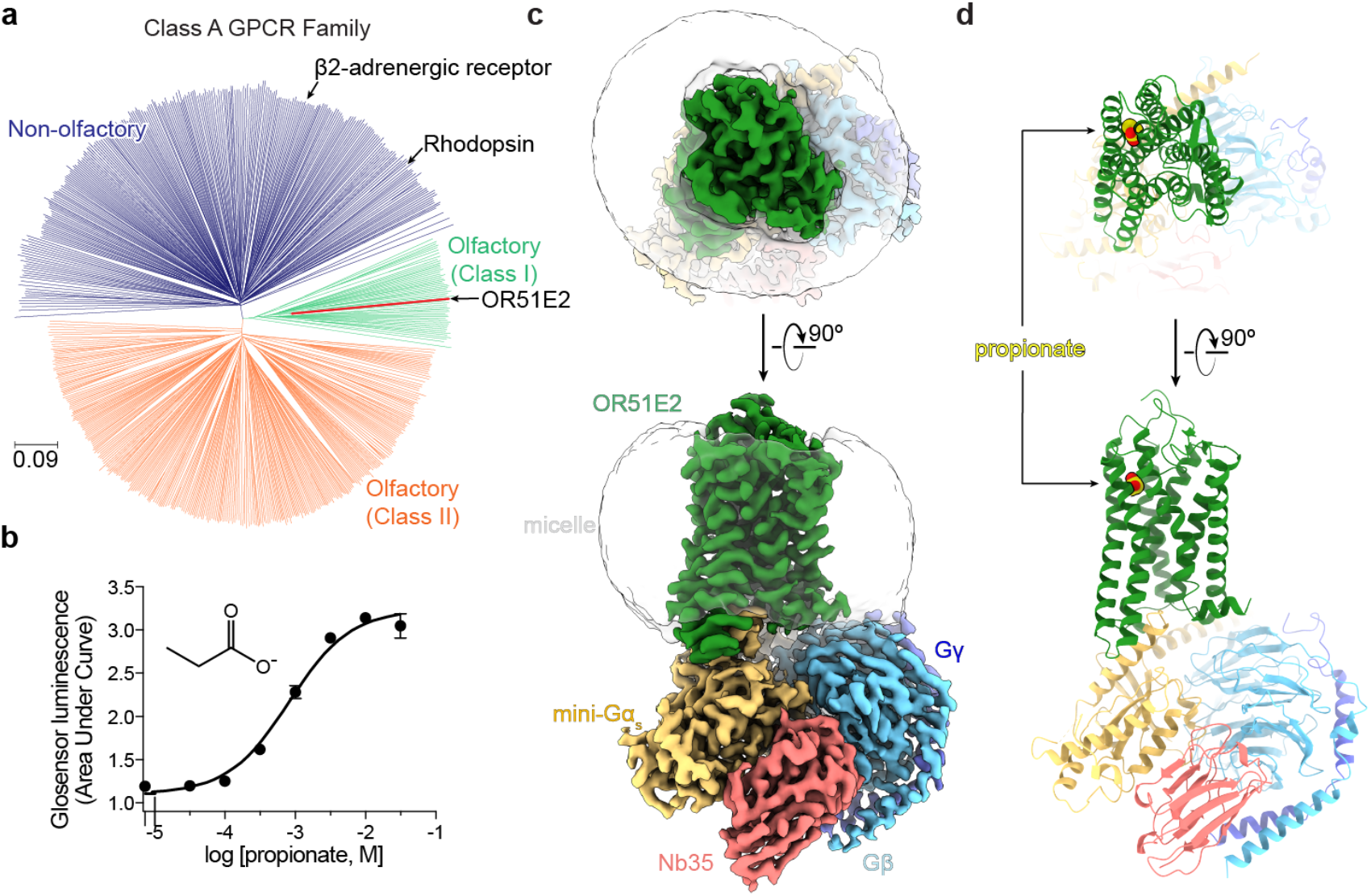
Structure of human olfactory receptor OR51E2. **a)** Phylogenetic tree of human Class A GPCRs, including both non-olfactory (blue) and olfactory receptors. Olfactory receptors are further divided into Class I (green) and Class II (orange). OR51E2 is a Class I OR. The phylogenetic distance scale is represented on the left bottom corner (the distance represents 9% differences between sequences). **b)** Real-time monitoring of cAMP concentration assay showing that human OR51E2 responds to the odorant propionate. Data points are mean ± standard deviation from *n* = 4 replicates. Cryo-EM density map (**c**) and ribbon model (**d**) of active human OR51E2 bound to propionate (yellow spheres) in complex with G_s_ heterotrimer and stabilizing nanobody Nb35.

Odorant stimulation of ORs activates signaling pathways via the stimulatory G protein G_olf_, which ultimately leads to excitation of OSNs^8^. Each OR can only interact with a subset of all potential odorants. Conversely, a single odorant can activate multiple ORs^2^. This principle of molecular recognition enables a central neural logic of olfaction where the perception of smell arises from the combinatorial activity of multiple unique ORs that respond to an individual odorant^2^. Because each mature OSN expresses only a single OR gene^9^, understanding how an individual OR is activated provides direct insight into the sensory coding of olfaction.

To understand olfaction at a fundamental level, we need a structural framework describing how odorants are recognized by ORs. Although recent structures of insect odorant-gated ion channels have begun to decipher this molecular logic^10,11^, the molecular rules that govern odorant recognition in vertebrate ORs are likely distinct and remain obscure. Here, we used cryogenic electron microscopy (cryo-EM) to determine the structure of a human OR activated by an odorant. This structure reveals specific molecular interactions that govern odorant recognition and provides a foundation for understanding how odorant binding activates ORs to instigate cellular signaling.

### Structure of odorant bound OR51E2

Several challenges have limited structural interrogation of vertebrate ORs, including low expression levels in heterologous systems, low solubility of most volatile odorants, and precipitous instability of purified ORs^12–15^. We therefore sought to identify a human OR that overcomes these challenges. We prioritized a subset of ORs that are also expressed in tissues outside of OSNs with chemoreceptive functions that are independent of olfaction^16^. The ability of these ORs to function in non-olfactory tissue suggested that they may be more amenable to expression in heterologous cell expression systems that lack olfactory-tissue specific chaperones^13^. In a second line of reasoning, we prioritized Class I (so called “fish-like”) ORs as these receptors generally recognize water-soluble odorants^17^. By contrast, Class II ORs tend to respond to more hydrophobic odorants. Finally, we prioritized ORs that have significant conservation across evolution, potentially because they recognize odorants that are critical for animal survival across many species^5^. We reasoned that such ORs may be more constrained by evolution for stability. With this approach, we identified human OR51E2 as an ideal candidate for structure determination (**Supplementary Fig. 1**). OR51E2 is a Class I OR that responds to the short chain fatty acid propionate^18^ (**Fig. 1a,b**). In addition to its olfactory function, OR51E2 and its mouse homolog Olfr78 are expressed in several other tissues to enable chemoreception of short chain fatty acids^19–24^. Consistent with our reasoning, OR51E2 emerged as one of the most highly expressed ORs in HEK293T cells among hundreds of human and mouse ORs that we have previously tested^12^.

To further stabilize OR51E2, we aimed to isolate OR51E2 in a complex with a heterotrimeric G protein. ORs couple with the two highly homologous stimulatory G proteins Gα_olf_ and Gα_s_. In mature OSNs, ORs activate Gα_olf_ to stimulate cAMP production via adenylyl cyclase^8^. In immature OSNs, ORs activate adenylyl cyclase via Gα_s_ to drive accurate anterior-posterior axon targeting^25^. Furthermore, OR51E2 signals via Gα_s_ outside of the olfactory system in tissues lacking Gα_olf_^20^. The ability of OR51E2 to signal physiologically via Gα_s_, combined with the availability of a nanobody (Nb35) that stabilizes GPCR-Gα_s_ complexes^26^, prompted us to focus on purifying an OR51E2-G_s_ complex. To do so, we generated an OR51E2 construct with a C-terminally fused “miniGa_s_” protein. The miniGa_s_ protein is engineered to trap the receptor-interacting conformation of Ga_s_ in the absence of any guanine nucleotide^27^. Fusion of the miniG_s_ to OR51E2 fully blocked propionate stimulated cAMP signaling in HEK293T cells (**Supplementary Fig. 2b**). We surmised that miniGα_s_ tightly engages the 7TM core of OR51E2 to preclude endogenous Ga_s_ coupling and cAMP production.

We purified OR51E2-miniG_s_ in the presence of 30 mM propionate, and then further generated a complex with recombinantly purified Gβ_1_γ_2_ and Nb35 (**Supplementary Fig. 2a and c**). The resulting preparation was vitrified and analyzed by single particle cryogenic electron microscopy (cryo-EM) (**Supplementary Fig 3** and **Supplementary Table 1**), which yielded a 3.1 Å resolution map of OR51E2 bound to the G_s_ heterotrimer. We additionally generated a map with focused refinement on only the 7TM domain of OR51E2, which afforded improved map resolution of the binding site and extracellular loops of the receptor (**Supplementary Fig. 3e**). The resulting reconstructions allowed us to model the OR51E2 7TM domain, the propionate ligand, and the G_s_ heterotrimer (**Fig. 1c,d** and **Supplementary Fig. 4a-c**).

### Odorant binding pocket

We identified cryo-EM density for propionate in a region bounded by transmembrane helices (TM) 3, 4, 5, and 6 in OR51E2 (**Fig. 2a** and **Supplementary Fig. 4b,d**). The propionate odorant binding pocket in OR51E2 is in a similar general region as ligand binding pockets in two prototypical Class A GPCRs: the adrenaline binding site in the β2-adrenergic receptor (β2-AR)^28^ and all-trans retinal in rhodopsin^29^ (**Fig. 2a–c**). Compared to the β2-AR and rhodopsin, the odorant binding pocket in OR51E2 is smaller and does not engage TM2 and TM7. Extensive packing of the OR51E2 N-terminus with extracellular loops 1 and 2 (ECL1 and ECL2) diminishes the potential size of the odorant binding pocket. Notably, unlike many class A GPCRs with diffusible agonists, the binding pocket for propionate is fully occluded from the extracellular milieu (**Fig. 2d**).

**Figure 2.**
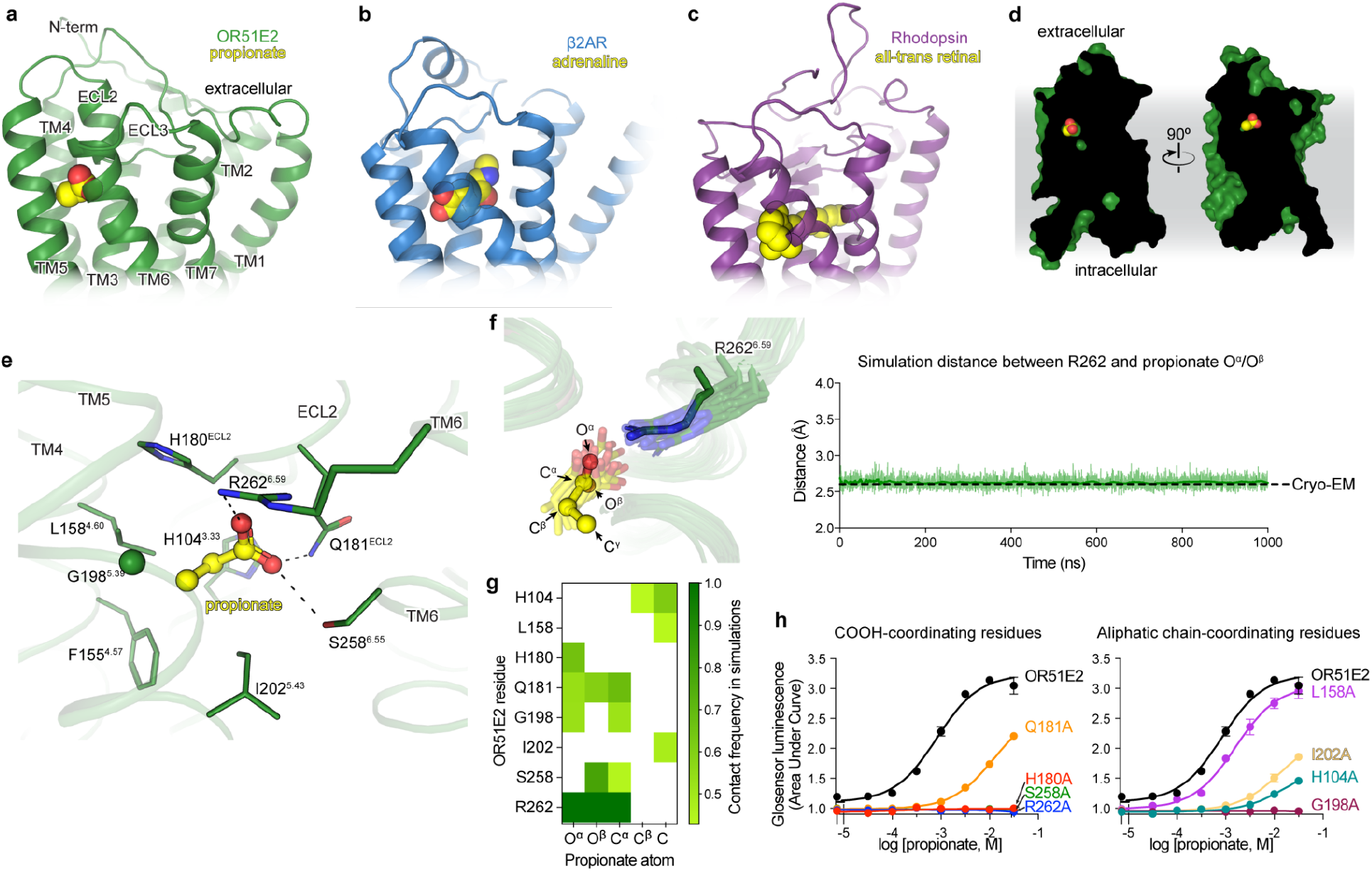
Odorant binding pocket in OR51E2. Comparison of propionate binding site in OR51E2 (**a**) to two other prototypical Class A GPCRs, the β2-adrenergic receptor (β2AR) bound to adrenaline (PDB 4LDO)^28^ (**b**) and rhodopsin bound to all-trans retinal (PDB 6FUF)^29^ (**c**). Propionate primarily contacts TM4, TM5, TM6 and ECL2. By contrast adrenaline and all-trans retinal make more extensive contacts with other GPCR transmembrane helices. **d**) The binding site of propionate in active OR51E2 is occluded from extracellular solvent. **e**) Close-up view of propionate binding site in OR51E2. **f**) Molecular dynamics simulations snapshots of OR51E2 bound to propionate are shown as transparent sticks and overlaid on the cryo-EM structure. R262^6.59^ makes a persistent contact with propionate over 1000 ns of simulation time (see Supplementary Fig. 8 for simulation replicates). The minimum distance between any of R262^6.59^ sidechain nitrogens and propionate oxygens is shown. **g**) Heatmap of contact frequencies of interaction between OR51E2 binding site residues and propionate atoms (as labeled in (f)) obtained from five independent molecular dynamics simulations each 1 μs long (total time 5 μs). Contact frequency cutoff between receptor residue and ligand atoms were set at 40%. **h**) Alanine mutagenesis analysis of propionate-contacting residues in OR51E2 using a real-time monitoring of cAMP concentration assay. Data points are mean ± standard deviation from *n* = 3 experiments.

Propionate makes several contacts within the OR51E2 odorant binding pocket. The carboxylic acid of propionate engages R262^6.59^ (superscript numbers indicate conserved Ballesteros-Weinstein numbering for GPCRs^30,31^) in TM6 as a counter-ion. The same propionate functional group also engages in hydrogen bonding interactions with S258^6.55^ and Q181 in ECL2 (**Fig. 2e**). We used molecular dynamics (MD) simulations to understand whether these interactions are stable. We performed five 1 μs simulations of OR51E2 bound to propionate, but in the absence of the G_s_ heterotrimer. During these simulations, we observed that the carboxylic group of propionate forms a persistent interaction with R262^6.59^, with an average distance that is identical to that observed in the cryo-EM structure (**Fig. 2f** and **Supplementary Fig. 5**). Simulations also supported persistent interactions between the propionate carboxylic group and S258^6.55^, with additional contacting residues outlined in **Fig 2g**. Indeed alanine mutations for these carboxylic group coordinating residues, with the exception of Q181^ECL2^, abolished propionate induced activation of OR51E2 (**Fig. 2h**).

The van-der Waals contacts between the propionate aliphatic group and OR51E2 are governed by tight packing interactions. The aliphatic portion of propionate contacts residues in TM3 (H104^3.33^), TM4 (F155^4.57^ and L158^4.60^), and TM5 (G198^5.39^ and I202^5.43^). Unlike the persistent contacts observed for the oxygens in the carboxylic acid group, interactions between specific propionate carbon atoms and aliphatic residues in OR51E2 were more dynamic in simulations (**Fig. 2g**) and showed minimal contact with F155^4.57^. However, alanine mutations to G198^5.39^, I202^5.43^ and H104^3.33^ significantly decreased propionate activity at OR51E2, suggesting that there are specific spatial requirements for propionate to bind and activate the receptor. By contrast, propionate is only moderately less efficacious at OR51E2 with the L158^4.60^A mutation (**Fig. 2h**), likely because this residue only engages the distal Cγ carbon of propionate. OR51E2 therefore recognizes propionate with specific ionic and hydrogen bonding interactions combined with more distributed van der Waals interactions with tight shape complementarity.

### Tuning olfactory receptor selectivity

Many ORs are capable of responding to a wide diversity of chemically distinct odorants^2,18^. Class I ORs, by contrast, are generally more restricted to carboxylic acid odorants^32^. We tested the selectivity of fatty acid odorants of various chain lengths at OR51E2 to understand how structural features in the receptor lead to odorant specificity. Consistent with previous reports^23,33^, we identified that acetate (C2) and propionate (C3) activate OR51E2 with millimolar potency (**Fig. 3a,b**). By contrast, longer chain length fatty acids (C4-C10) were either poorly or not active at OR51E2.

**Figure 3.**
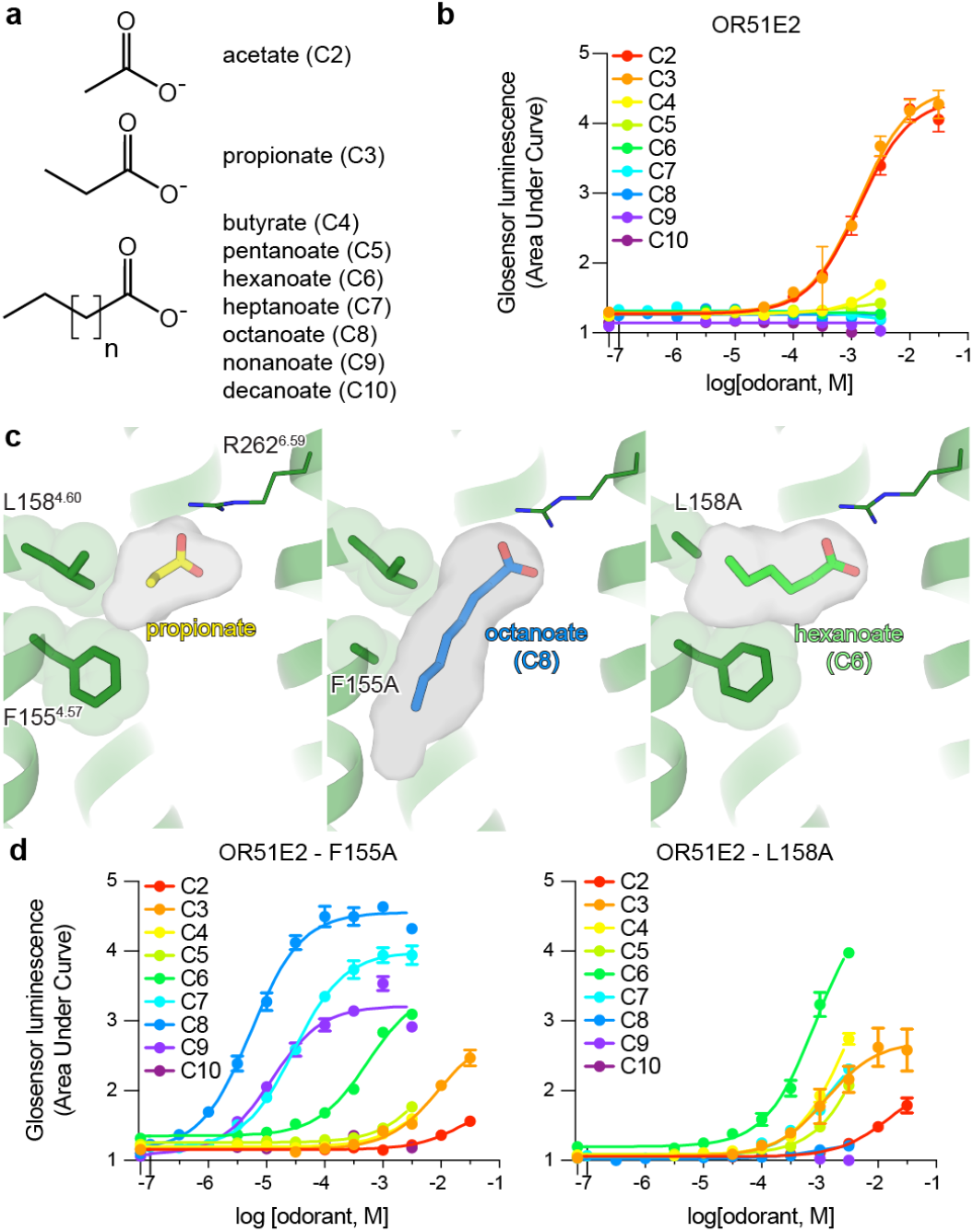
Tuning OR51E2 odorant selectivity. **a,b)** OR51E2 responds selectively to the short chain fatty acids acetate and propionate as measured by a cAMP production assay. **c)** Close up view of the OR51E2 binding pocket with the binding pocket cavity shown as gray surface. Replacement of F155^4.57^ and L158^4.60^ with alanine is predicted to yield a binding pocket with increased volume capable of accommodating a longer chain fatty acid. Docked poses of octanoate (C8) and hexanoate (C6) are shown in the F155A and L158A mutants of OR51E2, respectively. **d)** The F155A and L158A mutations in OR51E2 lead to increased sensitivity to long chain fatty acids. Conversely, the potency for acetate and propionate is reduced for these two mutants. Data points in b and d are mean ± standard deviation from *n* = 4 experiments.

We speculated that the selectivity of OR51E2 for short chain fatty acids arises from the restricted volume of the occluded binding pocket (31 Å^3^), which would accommodate short chain fatty acids like acetate and propionate but would preclude binding of fatty acids with longer aliphatic chain lengths (**Fig. 3c**). We therefore hypothesized that the volume of the binding pocket acts as a selectivity determinant for fatty acid chain length. To directly test this hypothesis, we designed two mutations that are predicted to result in increased binding pocket volumes while maintaining the specific contacts with R262^6.59^ important for fatty acid activation of OR51E2. More specifically, we mutated two residues that are proximal to the carbon chain of propionate: F155^4.57^ and L158^4.60^. Computational modeling of the F155^4.57^A and L158^4.60^A mutations predicted pocket volumes of 90 Å^3^ and 68 Å^3^ respectively, suggesting that both mutants should sufficiently accommodate fatty acids with longer chain length (**Fig. 3c**). Indeed in cAMP assays, both the F155^4.57^A and L158^4.60^A OR51E2 mutants were broadly responsive to longer chain fatty acids (**Fig. 3d, Supplementary Table 2** and **3**). The size of each binding pocket was correlated with the maximum chain length tolerated and, additionally, which chain length has the greatest potency. For example, F155^4.57^A is responsive to a range of fatty acids (C2-C9), with octanoate (C8) displaying maximal potency and efficacy. By contrast, hexanoate (C6) is the most efficacious agonist at the L158^4.60^A mutant. For both of these mutations, the potency of acetate and propionate is reduced compared to OR51E2, suggesting that tight packing interactions with the aliphatic chain is an important determinant of agonist potency.

We next examined the conservation of selectivity-determining residues in both human Class I and Class II ORs. Reflecting its importance in fatty acid recognition, arginine is highly conserved in the 6.59 position in most human Class I ORs (Class I 71% vs Class II 7%) (**Supplementary Fig. 6**). Positions 4.57 and 4.60 in all human Class I ORs are constrained to aliphatic amino acids of different size (V/I/L/M/F, Class I >80% vs Class II <15%). By contrast, none of these positions have similar constraints in Class II ORs. We surmise that the conserved residue R^6.59^ may anchor odorants in many Class I OR binding pockets, while diversity in the 4.57 and 4.60 positions tunes the binding pocket to enable selective recognition of the remainder of the molecule. Two features may therefore drive odorant recognition for Class I ORs: 1) hydrogen-bonding or ionic interactions that anchor polar features of odorants to conserved OR binding pocket residues, and 2) van-der Waals interactions of diverse aliphatic residues in the OR binding pocket that define a closed volume having a geometry that closely matches the shape of cognate odorants.

### Activation mechanisms of OR51E2

Odorant binding to ORs is predicted to cause conformational changes in the receptor that enable G protein engagement. Our strategy to stabilize OR51E2 with miniG_s_ precluded structure determination of inactive OR51E2 in the absence of an odorant. We therefore turned to comparative structural modeling, mutagenesis studies, and molecular dynamics simulations to understand the effect of propionate binding on the conformation of OR51E2.

Comparison of active OR51E2 to G_s_-coupled, active state β2-adrenergic receptor (β2-AR) demonstrated that both receptors engage the G protein with a similar overall orientation of the 7TM domain and Gα_s_ (**Fig. 4a** and **Supplementary Fig. 7**). A central hallmark of GPCR activation is an outward displacement of TM6 in the cytoplasmic side of the receptor, which is accompanied by more subtle movement of the other TM helices^34^. These conformational changes create a cavity for the G protein C-terminal α-helix. Prior structural biology studies have identified two regions conserved in Class A GPCRs that are critically important for allosteric communication between the agonist binding site and the G protein-binding site: a connector region that is adjacent to the ligand binding site and a G protein-coupling region adjacent to the Gα_s_ C-terminal α-helix^34^ (**Fig. 4a**). We aimed to understand how propionate binding to OR51E2 stabilizes these regions in an active conformation. Although the overall conformation of OR51E2 and β2-AR are similar (root-mean-square deviation (RMSD) of 3.2 Å), the specific sequences that define the G protein-coupling and connector regions are distinct between ORs and non-olfactory Class A GPCRs. Comparison of sequence conservation in TM6 between human ORs and non-olfactory Class A GPCRs revealed a highly conserved motif (KAFSTCxSH^6.40^) in the G protein coupling region in ORs that is absent in non-olfactory receptors (**Fig. 4b**). By contrast, the highly conserved CWxP^6.50^ motif in the connector region of Class A GPCRs is absent in ORs. Instead ORs contain the previously described FYGx^6.50^ motif in the connector region^35^ (**Fig. 4f,g**).

**Figure 4.**
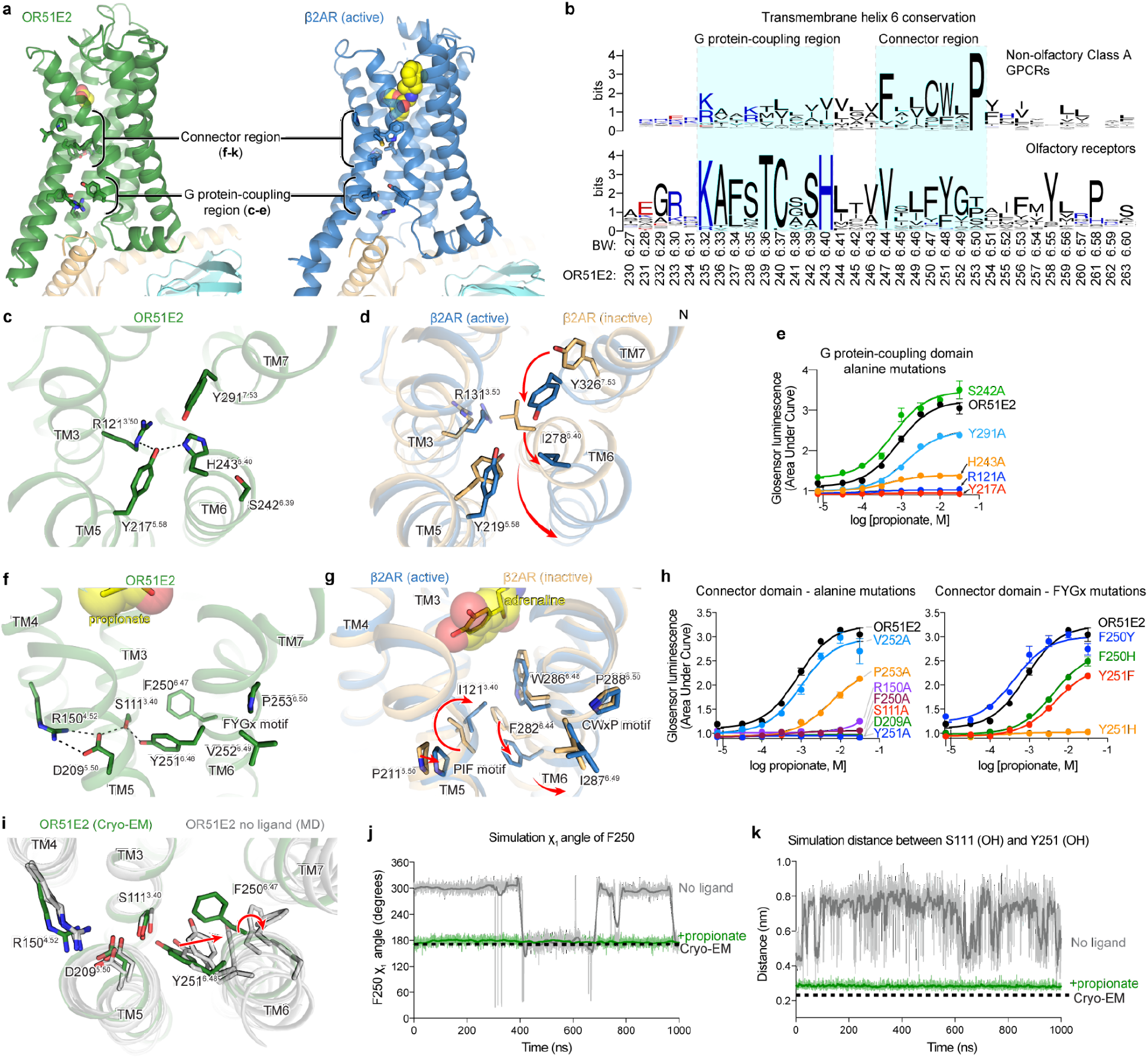
Activation mechanism of OR51E2. **a)** Ribbon diagram comparing structures of propionate-bound OR51E2-miniG_s_ complex (green) to BI-167107 bound β2AR-G_s_ complex (blue, PDB 3SN6). For both receptors, the connector region couples conformational changes at the ligand binding site with the G protein-coupling region. **b)** Weblogo depicting conservation of transmembrane helix 6 amino acids in either human olfactory receptors or human non-olfactory Class A GPCRs. Amino acid numbering for OR51E2 and Ballosteros-Weinsten (BW) are indicated. Close-up view of the G protein-coupling domain in active OR51E2 (**c)** and both active and inactive β2AR (**d**). Activation of β2AR is associated with an inward movement of TM7 and a contact between Y219^5.58^ and Y326^7.53^. In OR51E2, by contrast, H243^6.40^ interacts with Y217^5.58^ in the active state. **e**) Alanine mutagenesis analysis of G protein-coupling domain residues in OR51E2 using a real-time monitoring of cAMP concentration assay. Close-up views of the connector region in active OR51E2 (**f)** and both active and inactive β2AR (**g**). **h**) Mutagenesis analysis of connector region residues in OR51E2 using a real-time monitoring of cAMP concentration assay. **i**) Snapshots from molecular dynamics (MD) simulations of OR51E2 with propionate removed. Simulations show increased flexibility of TM6 in the connector region residues. Molecular dynamics trajectories for representative simulations showing rotation of side chain rotamer angle of F250^6.47^ (**j**) and distance between S111^3.40^ and Y251^6.48^ hydroxyl groups (**k**) performed with or without propionate over the course of 1000 ns MD simulation (see Supplementary Fig. 8 for simulation replicates). Data points in e and h mean ± standard deviation from *n* = 4 experiments.

Closer inspection of the G protein-coupling region in OR51E2 revealed a unique hydrogen-bonding network between the highly conserved residues R121^3.50^ in TM3, H243^6.40^ in TM6 and Y217^5.58^ in TM5 that is not observed in other Class A GPCRs (**Fig. 4c,d**). Activation of the β2-AR is associated with an inward movement of TM7 that positions Y316^7.53^ within a water-mediated hydrogen bonding distance of Y219^5.58^; this movement leads to outward movement of TM6 by displacing the aliphatic I278^6.40^ residue (**Fig. 4d**). Given the high conservation of R^3.50^, H^6.40^ and Y^5.58^ across all ORs (89%, 97% and 93%, respectively, **Supplementary Fig. 7**), we propose that this contact is important in stabilizing the OR active conformation. Indeed alanine mutagenesis of OR51E2 residues in the G protein coupling region show a dramatic loss of activity for H243^6.40^, Y217^5.58^, and R121^3.50^ associated with poor receptor expression (**Fig. 4e** and **Supplementary Table 2**). Mutation of Y291^7.53^ in OR51E2, by contrast, has a more modest effect on propionate activity.

We next examined the connector region of OR51E2 directly adjacent to the propionate binding site (**Fig. 4f**). Activation of the β2-AR is associated with a rearrangement of the PIF motif between positions I^3.40^ (TM3), P^5.50^ (TM5), and F^6.44^ (TM6), which leads to an outward displacement of TM6. This coordinated movement has been observed for the majority of class A GPCRs for which both active and inactive state structures have been obtained ^34^. Conservation at the PIF positions is low in ORs, suggesting an alternative mechanism. In OR51E2, we observe an extended hydrogen bonding network between Y251^6.48^ of the OR-specific FYGx motif and residues in TM3 (S111^3.40^), TM4 (R150^4.52^), and TM5 (D209^5.50^). Notably, the intramembrane ionic interaction between D209^5.50^ and R150^4.52^ is likely only conserved in Class I ORs (Class I: D^5.50^-82%, R^4.52^-88%, Class II: D^5.50^-0.3%, R^4.52^-0%, **Supplementary Fig. 7**). Alanine mutagenesis of most residues in this connector region of OR51E2 abolishes response to propionate (**Fig. 4h**), in part because mutations in this region dramatically decrease receptor expression (**Supplementary Table 2**). More conservative substitutions to F250^6.47^ or Y251^6.48^ also show impairment in OR51E2 function, suggesting that the specific contacts observed in active OR51E2 are important for robust receptor activation.

We turned to molecular dynamics simulations to examine how ligand binding influences the conformation of the connector region. After removing the G protein, we simulated OR51E2 with and without propionate in the binding site. For each condition, we performed five 1 μs simulations. OR51E2 simulated with propionate remains in a conformation similar to the cryo-EM structure. In the absence of propionate, the connector region of OR51E2 displays significantly more flexibility in simulations (**Fig. 4i** and **Supplementary Fig. 8**). We observed two motions in the FYGx motif associated with this increased conformational heterogeneity: a rotameric flexibility of F250^6.47^ between the experimentally observed conformation and alternative rotamers and a disruption of a hydrogen bond between Y251^6.48^ and S111^3.40^ (**Fig. 4j,k** and **Supplementary Fig. 8**). Simulations without propionate show that the distance between the hydroxyl groups of Y251^6.48^ and S111^3.40^ is >4 Å, indicating the loss of a hydrogen bond that was observed in both the cryo-EM structure of OR51E2 and the MD simulations with propionate (**Fig. 4k** and **Supplementary Fig. 8**). Based on structural comparison to other Class A GPCRs, mutagenesis studies, and molecular dynamics simulations, we therefore propose that odorant binding stabilizes the conformation of an otherwise dynamic FYGx motif to drive OR activation.

### Structural dynamics of ECL3 in OR function

Olfactory receptors display significant sequence variation in extracellular loop 3 (ECL3), a region previously shown to be critical for recognition of highly diverse odorants^36,37^. We therefore aimed to understand the involvement of ECL3 in propionate binding to OR51E2, and more generally, how ECL3 may drive the conformational changes in TM6 necessary for OR activation (**Fig. 5**). In our structure of OR51E2, ECL3 is directly coupled to odorant binding via a direct interaction between the carboxylic acid moiety of propionate and the ECL3 adjacent residue R262^6.59^ (**Fig. 5a**). In order to investigate the role of R262^6.59^ in maintaining the conformation of ECL3 by binding the odorant, we analyzed simulations of OR51E2 performed without propionate. In the absence of coordination with the carboxylic acid group of propionate, R262^6.59^ showed a marked increase in flexibility, with an outward movement of up to 8 Å away from the ligand binding site (**Fig. 5b** and **c**). This movement is accompanied by displacement of ECL3 away from the odorant binding pocket.

**Figure 5:**
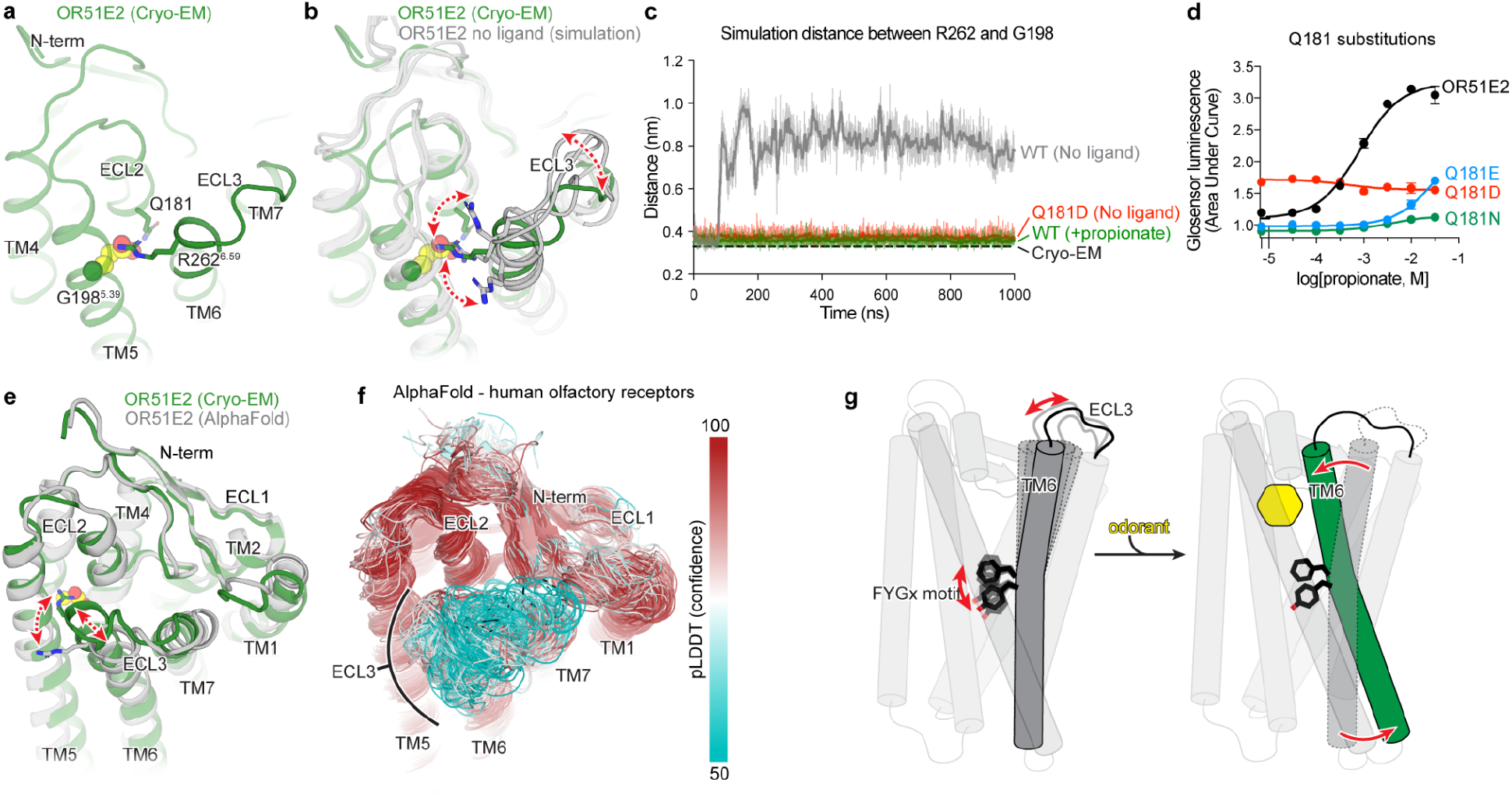
Structural dynamics of ECL3 in OR function. **a)** Residue R262^6.59^ in ECL3 makes a critical contact with propionate. Residue Q181^ECL2^ in ECL2 is highlighted. **b)** Molecular dynamics snapshots of OR51E2 simulated in the absence of propionate shows increased flexibility of R262^6.59^. **c)** In simulations of wild-type (WT) OR51E2 bound to propionate, the distance between R262^6.59^ and G198^5.39^ is stable and similar to the cryo-EM structure. Simulations of WT OR51E2 without propionate (no ligand) show increased distances between R262^6.59^ and G198^5.39^. In simulations of Q181D mutant without propionate, the distance between R262^6.59^ and G198^5.39^ is similar to WT OR51E2 bound to propionate. Distance was measured between R262^6.59^ sidechain atoms and G198^5.39^ main chain atoms (excluding the hydrogens) over the course of 1000 ns MD simulation (see Supplementary Fig. 8 for simulation replicates). **d)** Conservative mutagenesis of Q181^ECL2^ shows that the Q181D mutant is constitutively active, potentially because it substitutes a carboxylic acid in the OR51E2 binding pocket. **e)** Comparison of cryo-EM structure of OR51E2 with the AlphaFold2 predicted structure shows high similarity in the extracellular domain with the exception of the ECL3 region. The AlphaFold2 model shows an outward displacement of R262^6.59^ and ECL3 similar to simulations of apo OR51E2. **f)** AlphaFold2 predictions for all human olfactory receptors show low confidence in the ECL3 region and high confidence in other extracellular loops. **g)** A model for ECL3 as a key site for olfactory receptor activation.

To test whether inward movement of R262^6.59^ is itself sufficient to activate OR51E2, we designed a gain-of-function experiment. We hypothesized that introduction of an acidic residue in the binding pocket with an appropriate geometry may substitute for the carboxylic acid of propionate and coordinate R262^6.59^. Indeed, substitution of Asp in position 181 (Q181D) of OR51E2 yielded increased cAMP basal activity (**Fig. 5d**). By contrast, introduction of Glu in the same position (Q181E) rendered OR51E2 largely inactive, suggesting the requirement for a precise coordination geometry for R262^6.59^. Substitution with the larger Gln (Q181N) rendered OR51E2 completely unresponsive to propionate, either by sterically blocking R262^6.59^ or by displacing propionate itself. In simulations of OR51E2 with the Q181D substitution, R262^6.59^ is persistently engaged toward the ligand binding site (**Fig. 5b**). Furthermore, this inward movement of R262^6.59^ and ECL3 is accompanied by activation-associated conformational changes in the connector domain of OR51E2 (**Supplementary Fig. 9**), perhaps explaining the basal activity of Q181D mutant. Inward movement of ECL3 is therefore sufficient to activate OR51E2.

Because conformational changes in ECL3 are critical to OR51E2 activation, we speculated that this region may provide a common activation mechanism across the OR family. To probe this notion, we examined structural predictions of all human olfactory receptors by AlphaFold2^38^. We first compared the AlphaFold2 prediction for OR51E2 with the cryo-EM structure, which yielded a high degree of agreement reflected in a RMSD of 1.3 Å for Cα atoms. Importantly, the AlphaFold2 predicted structure of OR51E2 appears to be in an intermediate or inactive conformation characterized by outward displacement of R262^6.59^ and ECL3, a G protein-coupling domain in the inactive conformation, and TM6 more inwardly posed compared to active OR51E2 (**Fig 5e** and **Supplementary Fig. 10**). We next examined the predicted structures of all human ORs, which revealed a largely shared topology for the extracellular region for the broader family (**Fig. 5f**). Indeed, the per-residue confidence score from AlphaFold2 (predicted local distance difference test, pLDDT) for the N terminus, ECL1, and ECL2 are predicted with high confidence for the most ORs. By contrast, ECL3 shows significantly lower pLDDT scores. Because low pLDDT scores correlate with disordered protein regions^38^, we surmise that, in the absence of odorant binding, the structure of ECL3 is less constrained compared to the rest of the odorant binding pocket for the broader OR family. Similar to OR51E2, odorant binding may therefore stabilize ECL3 to drive receptor activation for the broader OR family.

## Discussion

We propose the following model for OR51E2 activation (**Fig. 5g**). In the unbound state, the extracellular segment of TM6 is dynamic. Upon binding of propionate, TM6 rotates inward towards the 7TM domain and is stabilized via a direct coordination of the propionate carboxylic acid via R262^6.59^. The conserved FYGx motif in TM6 acts as a structural pivot point around which TM6 rotates to displace the intracellular end of TM6 from the TM-core and open the canonical active G protein-binding site. Although specific interactions between the propionate aliphatic chain and residues within the binding site are important for achieving full potency of the odorant response, OR51E2 is constitutively active when an aspartate residue (Q181D) is introduced in the binding pocket. This suggests that the observed rotation of TM6 mediated by coordination of R262 with a stable anionic group in the binding site, in itself is sufficient for receptor activation. While this model remains speculative due to the lack of an experimentally-determined inactive-state structure of OR51E2, it integrates the findings from unique structural features of ORs compared to other Class A GPCRs, molecular dynamics simulations, and mutagenesis studies. A similar mechanism may be responsible for the activation of most Class I ORs, a large majority of which recognize carboxylic acids and contain an arginine at position 6.59. The mechanism of activation of Class II ORs, which recognize a broader range of volatile odorants and lack R^6.59^, could be potentially distinct.

Our work illuminates the molecular underpinnings of odorant recognition in a vertebrate Class I OR. While the full breadth of potential odorants that activate OR51E2 remains to be characterized, profiling of known fatty acid odorants suggests that OR51E2 is narrowly tuned to short chain fatty acids^18,23^. Propionate binds OR51E2 with two types of interactions - specific ionic and hydrogen bonding interactions that anchor the carboxylic acid, and more nonspecific hydrophobic contacts that rely on shape complementarity with the aliphatic portion of the ligand. We demonstrate that the specific geometric constraints imposed by the occluded OR51E2 odorant binding pocket are responsible, in part, for this selectivity. Molecular recognition in OR51E2 is therefore distinct from the distributed hydrophobic interactions that mediate odorant recognition at an insect odorant-gated ion channel^11^. We anticipate that the molecular mechanism we define here for OR51E2 is likely to extend to other Class I ORs that recognize polar, water soluble odorants with multiple hydrogen bond acceptors and donors. Molecular recognition by more broadly tuned ORs, and the larger Class II OR family, however, remains to be defined.

The structural basis of ligand recognition for OR51E2 also provides insight into evolution of the OR family. Unlike most vertebrate OR genes that have evolved rapidly via gene duplication and diversification, OR51E2 is one of a few ORs with strong evolutionary conservation within different species^5^. This constraint may result from recognition of odorants important for survival or from vital non-olfactory roles of OR51E2 activity detecting propionate and acetate, the main metabolites produced by the gut microbiota. Molecular recognition of propionate by OR51E2 may therefore represent a unique example of specificity within the broader OR family. While future work will continue to decipher how hundreds of ORs sense an immensely large diversity of odorants, our structure and mechanistic insight into OR51E2 function provides a new foundation to understand our sense of smell at an atomic level.

## METHODS

### Expression and purification of OR51E2-miniG_s_ protein

Human *OR51E2* was cloned into pCDNA-Zeo-TetO, a custom pcDNA3.1 vector containing a tetracycline inducible gene-expression cassette^41^. The construct included an N-terminal influenza hemagglutinin signal sequence and the FLAG (DYKDDDK) epitope tag. The construct further included the miniG_s399_ protein^5^, which was fused to the C-terminus of OR51E2 with a human rhinovirus 3C (HRV 3C) protease cleavage sequence flanked by Gly-Ser linkers.

The resulting construct (OR51E2-miniG_s399_) was transfected into 1 L of inducible Expi293F-TetR cells (unauthenticated and untested for mycoplasma contamination, Thermo Fisher) using the ExpiFectamine 293 Transfection Kit (Thermo Fisher) as per manufacturer’s instructions. After 16 hours, protein expression was induced with 1 μg/mL doxycycline hyclate (Sigma Aldrich), and the culture was placed in a shaking incubator maintaining 37°C and a 5% CO_2_ atmosphere. After 36 hours cells were harvested by centrifugation and stored at −80°C.

For receptor purification, cells were thawed and hypotonically lysed in 50 mM HEPES, pH 7.50, 1 mM EDTA, 30 mM sodium propionate (Sigma Aldrich), 100 μM tris(2-carboxyethyl)phosphine (TCEP, Fischer Scientific), 160 μg/mL benzamidine, 2 μg/mL leupeptin for 15 minutes at 4°C. Lysed cells were harvested by centrifugation at 16,000 x g for 15 min, and immediately dounce-homogenized in ice-cold solubilization buffer comprising 50 mM HEPES, pH 7.50, 300 mM NaCl, 1% (w/v) Lauryl Maltose Neopentyl Glycol (L-MNG, Anatrace), 0.1% (w/v) cholesteryl hemisuccinate (CHS, Steraloids), 30 mM sodium propionate, 5 mM adenosine 5’-triphosphate (ATP, Fischer Scientific), 2 mM MgCl2, 100 μM TCEP, 160 μg/mL benzamidine, and 2 μg/mL leupeptin. The sample was stirred for 2 hours at 4°C, and the detergent-solubilized fraction was clarified by centrifugation at 20,000 x g for 30 min. The detergent-solubilized sample was supplemented with 4 mM CaCl_2_ and incubated in batch with homemade M1-FLAG-antibody conjugated CNBr-sepharose under slow rotation for 1.5 hours at 4°C. The sepharose resin was transferred to a glass column and washed with 20 column volumes ice-cold buffer comprising 50 mM HEPES, pH 7.50, 300 mM NaCl, 0.05% (w/v) L-MNG, 0.005% (w/v) CHS, 30 mM sodium propionate, 2.5 mM ATP, 4 mM CaCl_2_, 2 mM MgCl2, and 100 μM TCEP. This was followed by 10 column volumes of ice-cold 50 mM HEPES, pH 7.50, 150 mM NaCl, 0.0075% (w/v) L-MNG, 0.0025% glyco-diosgenin (GDN, Anatrace), 0.001% (w/v) CHS, 30 mM sodium propionate, 4 mM CaCl_2_, and 100 μM TCEP. Receptor containing fractions were eluted with ice-cold 50 mM HEPES, pH 7.50, 150 mM NaCl, 0.0075% (w/v) L-MNG, 0.0025% (w/v) GDN, 0.001% (w/v) CHS, 30 mM sodium propionate, 5 mM EDTA, 100 μM TCEP, and 0.2 mg/mL FLAG peptide. Fractions containing OR51E2-miniG_s399_ fusion protein were concentrated in a 50 kDa MWCO spin filter (Amicon) and further purified over a Superdex 200 Increase 10/300 GL (Cytiva) size-exclusion chromatography (SEC) column, which was equilibrated with 20 mM HEPES, pH 7.50, 150 mM NaCl, 0.005% (w/v) GDN, and 0.0005% CHS, 30 mM sodium propionate, and 100 μM TCEP. Fractions containing monodisperse OR51E2-miniG_s399_ were combined and concentrated in a 50 kDa MWCO spin filter prior to complexing with Gβ_1_γ_2_ and Nb35.

### Expression and purification of Gβ_1_γ_2_

A baculovirus was generated with the pVLDual expression vector encoding both the human Gβ_1_ subunit with a HRV 3C cleavable N-terminal 6x His-tag and the untagged human Gγ_2_ subunit, in *Spodoptera frugiperda* Sf9 insect cells (unauthenticated and untested for mycoplasma contamination, Expression Systems). For expression, *Trichoplusia ni* Hi5 insect cells (unauthenticated and untested for mycoplasma contamination, Expression Systems) were infected at a density of 3.0 x 10^6^ cells/mL with high titer Gβ_1_γ_2_-baculovirus, and grown at 27 °C with 130 rpm shaking. After 48 hours, cells were harvested and resuspended in lysis buffer comprising 20 mM HEPES, pH 8.00, 5 mM β-mercaptoethanol (β-ME), 20 μg/mL leupeptin, and 160 μg/mL benzamidine. Lysed cells were pelleted at 20,000 x g for 15 min, and solubilized with 20 mM HEPES pH 8.0, 100 mM sodium chloride, 1% (w/v) sodium cholate (Sigma Aldrich), 0.05% (w/v) n-dodecyl-β-D-maltopyranoside (DM, Anatrace), and 5 mM β-mercaptoethanol (β-ME). Solubilized Gβ_1_γ_2_ was clarified by centrifugation at 20,000 x g for 30 min, and was then incubated in batch with HisPur Ni-NTA resin (Thermo Scientific). Resin-bound Gβ_1_γ_2_ was washed extensively, before detergent was slowly exchanged on-column to 0.1% (w/v) L-MNG, and 0.01% (w/v) CHS. Gβ_1_γ_2_ was eluted with 20 mM HEPES pH 7.50, 100 mM NaCl, 0.1% (w/v) L-MNG, 0.01% (w/v) CHS, 300 mM imidazole, 1 mM DL-Dithiothreitol (DTT), 20 μg/mL leupeptin, and 160 μg/mL benzamidine. Fractions containing Gβ_1_γ_2_ were pooled and supplemented with homemade 3C protease before overnight dialysis into buffer comprised of 20 mM HEPES pH 7.50, 100 mM NaCl, 0.02% (w/v) L-MNG, 0.002% (w/v) CHS, 1 mM DTT, and 10 mM imidazole. Uncleaved Gβ_1_γ_2_ was removed by batch incubation with Ni-NTA resin, before the unbound fraction containing cleaved Gβ_1_γ_2_ was dephosphorylated by treatment with lambda phosphatase (New England Biolabs), calf intestinal phosphatase (New England Biolabs), and antarctic phosphatase (New England Biolabs) for 1 hour at 4°C. Geranylgeranylated Gβ_1_γ_2_ heterodimer was isolated by anion exchange chromatography using a MonoQ 4.6/100 PE (Cytiva) column, before overnight dialysis in 20 mM HEPES, pH 7.50, 100 mM NaCl, 0.02% (w/v) L-MNG, and 100 μM TCEP. Final sample was concentrated on a 3 kDa MWCO spin filter (Amicon), and 20% (v/v) glycerol was added before flash freezing in liquid N_2_ for storage at −80°C.

### Expression and purification of Nb35

DNA encoding Nb35 (described by Rasmussen *et al.^6^*) was cloned into a modified pET-26b expression vector harboring a C-terminal His-tag followed by a Protein C (EDQVDPRLIDGK) affinity tag. The resulting DNA was transformed into competent Rosetta2 (DE3) pLysS *Escherichia coli* (UC Berkeley QB3 MacroLab) and inoculated into 100 ml Luria Broth supplemented with 50 μg/mL kanamycin, which was cultured overnight with 220 rpm shaking at 37°C. The following day, the starter culture was inoculated into 8 x 1 L of Terrific Broth supplemented with 0.1% (w/v) dextrose, 2 mM MgCl2, and 50 μg/mL kanamycin which were further cultured at 37°C with shaking. Nb35 expression was induced at OD_600_ = 0.6, by addition of 400 μM Isopropyl β-D-1-thiogalactopyranoside (IPTG, GoldBio) and lowering the incubator temperature to 20°C. After 21 hours of expression, cells were harvested by centrifugation and were resuspended in SET Buffer comprising 200 mM tris(hydroxymethyl)aminomethane (Tris, Sigma Aldrich), pH 8.00, 500 mM sucrose, 0.5 mM EDTA, 20 μg/mL leupeptin, 160 μg/mL benzamidine, and 1 U benzonase. After 30 minutes of stirring at RT, hypotonic lysis was initiated by a 3-fold dilution with deionized water. Following 30 minutes of stirring at RT, ionic strength was adjusted to 150 mM NaCl, 2 mM CaCl_2_, and 2 mM MgCl_2_ and the lysate was cleared by centrifugation at 20,000 x g for 30 min. The cleared lysate was incubated in batch with homemade anti-Protein C antibody coupled CNBr-sepharose under slow rotation. The resin was extensively washed with buffer comprising 20 mM HEPES, pH 7.50, 300 mM NaCl, and 2 mM CaCl_2_, and Nb35 was eluted with 20 mM HEPES pH 7.50, 100 mM NaCl, 0.2 mg/mL Protein C peptide, and 5 mM EDTA. Nb35 containing fractions were concentrated in a 10 kDa MWCO spin filter (Amicon) and further purified over a Superdex S75 Increase 10/300 GL column (Cytiva) SEC column equilibrated with 20 mM HEPES, pH 7.50, and 100 mM NaCl. Fractions containing monodisperse Nb35 were concentrated and supplemented with 20% glycerol prior to flash freezing in liquid N_2_ for storage at −80°C.

### Preparation of the active-state OR51E2-G_s_ complex

To prepare the OR51E2-G_s_ complex, a 2-fold molar excess of purified Gβ_1_γ_2_ and Nb35 was added to the SEC purified OR51E2-miniG_s399_ followed by overnight incubation on ice. The sample was concentrated on a 50 kDa MWCO spin filter (Amicon), and injected onto a Superdex 200 Increase 10/300 GL SEC column, equilibrated with 20 mM HEPES, pH 7.50, 150 mM NaCl, 0.0075% (w/v) L-MNG, 0.0025% (w/v) GDN, 0.001% (w/v) CHS, and 30 mM sodium propionate. Fractions containing the monomeric OR51E2-G_s_ complex were concentrated on a 100 kDa MWCO spin filter immediately prior to cryo-EM grid preparation.

### Cryo-EM vitrification, data collection, and processing

2.75 μL of purified OR51E2-G_s_ complex was applied to glow discharged 300 mesh R1.2/1.3 UltrAuFoil Holey gold support films (Quantifoil). Support films were plunge-frozen in liquid ethane using a Vitrobot Mark IV (Thermo Fisher) with a 10 s hold period, blot force of 0, and blotting time varying between 1-5 s while maintaining 100% humidity and 4°C. Vitrified grids were clipped with Autogrid sample carrier assemblies (Thermo Fisher) immediately prior to imaging. Movies of OR51E2-G_s_ embedded in ice were recorded using a Titan Krios Gi3 (Thermo Fisher) with a BioQuantum Energy Filter (Gatan) and a K3 Direct Electron Detector (Gatan). Data were collected using Serial EM^42^ running a 3 x 3 image shift pattern at 0° stage tilt. A nominal magnification of 105,000 x with a 100 μm objective was used in super-resolution mode with a physical pixel size of 0.81 Å pixel^-1^. Movies were recorded using dose fractionated illumination with a total exposure of 50 e^-^ Å^-2^ over 60 frames yielding 0.833 e^-^ Å^-2^ frames^-1^.

16,113 super-resolution movies were motion-corrected and Fourier cropped to physical pixel size using UCSF MotionCor2^43^. Dose-weighted micrographs were imported into cryoSPARC v3.2 (Structura Biotechnology), and contrast transfer functions were calculated using the Patch CTF Estimation tool. A threshold of CTF fit resolution > 5 Å was used to exclude low quality micrographs. Particles were template picked using a 20 Å low-pass filtered model that was generated *ab initio* from data collected during an earlier 200 kV screening session. 8,884,130 particles were extracted with a box size of 288 pixels binned to 72 pixels and sorted with the Heterogeneous Refinement tool, which served as 3D classification with alignment. Template volumes for each of the four classes were low-pass filtered to 20 Å and comprised an initial OR51E2-G_s_ volume as well as three scrambled volumes obtained by terminating the Ab-Initio Reconstruction tool before the first iteration. The resulting 1,445,818 particles were re-extracted with a box size of 288 pixels binned to 144 pixels and sorted by an additional round of Heterogeneous Refinement using two identical initial models and two scrambled models. 776,527 particles from the highest resolution reconstruction were extracted with an unbinned box size of 288 pixels, and were subjected to Homogeneous Refinement followed by Non-Uniform Refinement. Particles were exported using csparc2star.py from the pyem v0.5 script package^39^, and an inclusion mask covering the 7TM domain of OR51E2 was generated using the Segger tool in UCSF ChimeraX^44^ and the mask.py tool in pyem v0.5. Particles and mask were imported into Relion v3.0^45^ and sorted by several rounds of 3D classification without image alignment, where the number of classes and tau factor were allowed to vary. The resulting 204,438 particles were brought back into cryoSPARC and subjected to Non-Uniform Refinement. Finally, Local Refinement using an inclusion mask covering the 7TM domain was performed, using poses/shift gaussian priors with S.D. of rotational and shift magnitudes limited to 3° and 2 Å respectively.

### Model building and refinement

Model building and refinement were carried out using an Alphafold2^38^ predicted structure as a starting model, which was fitted into the OR51E2-G_s_ map using UCSF ChimeraX. A draft model was generated using ISOLDE^46^ and was further refined by iterations of real space refinement in Phenix^47^ and manual refinement in Coot^48^. The propionate model and rotamer library were generated with the PRODRG server^49^, docked using Coot, and refined in Phenix. Final map model validations were carried out using Molprobity and EMRinger in Phenix.

### Site Directed Mutagenesis

Generation of OR51E2 mutants was performed as described previously^50^. Forward and reverse primers coding for the mutation of interest were obtained from Integrated DNA Technologies. Two successive rounds of PCR using Phusion polymerase (Thermo Fisher Scientific: F-549L) were performed to amplify ORs with mutations. The first round of PCR generated two fragments, one containing the 5’ region upstream of the mutation site and the other the 3’ downstream region. The second PCR amplification joined these two fragments to produce a full ORF of the olfactory receptor. PCR products with desired length were gel purified and cloned into the MluI and NotI sites of mammalian expression vector pCI (Promega) that contains rho-tag. Plasmids were purified using the Thomas Scientific (1158P42) miniprep kit with modified protocol including phenol-chloroform extraction before column purification.

### cAMP signaling assays

The GloSensor cAMP assay (Promega) was used to determine real-time cAMP levels downstream of OR activation in HEK293T cells, as previously described^51^. HEK293T cells were cultured in Minimum Essential Media (MEM, Corning) supplemented by 10 % Fetal Bovine Serum (FBS,Gibco), 0.5 % Penicillin-Streptomycin (Gibco) and 0.5 % Amphotericin B (Gibco).

Cultured HEK293T cells were plated the day before transfection at 1/10 of 100 % confluence from a 100 mm plate into 96-well plates coated with poly D lysine (Corning). For each 96-well plate, 10 μg pGloSensor-20F plasmid (Promega) and 75 μg of Rho-tagged OR in the pCI mammalian expression vector (Promega) were transfected 18 to 24 h before odorant stimulation using Lipofectamine 2000 (Invitrogen: 11668019) in MEM supplemented by 10% FBS. On stimulation day, plates were injected with 25 μl of GloSensor substrate (Promega) and incubated for 2 hours in the dark at room temperature and in a odor-free environment. Odorants were diluted to the desired concentration in CD293 media (Gibco) supplemented with copper (30 μM CuCl_2_, Sigma-Aldrich) and 2 mM L-glutamine (Gibco) and pH adjusted to 7.0 with a 150 mM solution of sodium hydroxide (Sigma-Aldrich). After injecting 25 μl of odorants in CD293 media into each well, GloSensor luminescence was immediately recorded for 20 cycles of monitoring over a total period of 30 minutes using a BMG Labtech POLARStar Optima plate reader. The resulting luminescence activity was normalized to a vector control lacking any OR, and the OR response was obtained by summing the response from all 20 cycles to determine an area under the curve (AUC). Dose-dependent responses of ORs were analyzed by fitting a least squares function to the data using GraphPrism 9.

### Evaluating Cell Surface Expression

Flow-cytometry was used to evaluate cell surface expression of olfactory receptors as described previously^52^. HEK293T cells were seeded onto 35-mm plates (Greiner Bio-One) with approximately 3.5 x 10^5^ cells (25 % confluency). The cells were cultured overnight. After 18 to 24 hours, 1200 ng of ORs tagged with the first 20 amino acids of human rhodopsin (rho-tag) at the N-terminal ends^53^ in pCI (Promega) and 30 ng eGFP were transfected using Lipofectamine 2000 (Invitrogen: 11668019). 18 to 24 hours after transfection, the cells were detached and resuspended using Cell stripper (Corning) and then transferred into 5 mL round bottom polystyrene (PS) tubes (Falcon) on ice. The cells were spun down at 4°C and resuspended in phosphate buffered saline (PBS, Gibco) containing 15 mM NaN3 (Sigma-Aldrich) and 2% FBS (Gibco). They were stained with 1/400 (v/v) of primary antibody mouse anti rhodopsin clone 4D2 (Sigma-Aldrich: MABN15) and allowed to incubate for 30 minutes then washed with PBS containing 15 mM NaN3 and 2% FBS. The cells were spun again and then stained with 1/200 (v/v) of the phycoerythrin (PE)-conjugated donkey anti-mouse F(ab’)2 fragment antibody (Jackson Immunologicals: 715-116-150) and allowed to incubate for 30 minutes in the dark. To label dead cells, 1/500 (v/v) of 7-Amino-actinomycin D (Calbiochem: 129935) was added. The cells were then immediately analyzed using a BD FACSCanto II flow cytometer with gating allowing for GFP positive, single, spherical, viable cells and the measured PE fluorescence intensities were analyzed and visualized using Flowjo v10.8.1. Normalizing the cell surface expression levels of the OR51E2 mutants was performed using wild-type OR51E2 which showed robust cell surface expression and empty plasmid pCI which demonstrated no detectable cell surface expression.

### Molecular dynamics simulations

All MD simulations were performed using the GROMACS package^54^ (version 2021) with the CHARMM36m forcefield^55^ starting from the OR51E2 EM structure with and without propionate. The G protein was removed in all these simulations. The GPCR structures were prepared by Maestro “protein preparation wizard” module^56^. The missing side chains and hydrogen atoms were added. Furthermore, protein chain termini were capped with neutral acetyl and methylamide groups, and histidine protonated states were assigned, after which minimization was performed. The simulation box was created using CHARMM-GUI^57^. We used the PPM 2.0 function of OPM (Orientation of proteins in membranes) structure of OR51E2 for alignment of the transmembrane helices of protein structure and inserted into a 75% palmitoyl-oleoyl-phosphatidylcholine (POPC) / 25% Cholesteryl Hemi Succinate deprotonated (CHSD) bilayer. The CHSDs were placed around the GPCR structure. TIP3P water model was used for solvation and 0.15 M potassium chloride ions were added for neutralization. The final system dimensions were about 85 × 85 × 115 Å. The system was minimized with position restraints (10 kcal/mol/Å^2^) on all heavy atoms of GPCR and ligand, followed by a 1 ns heating step which raise the temperature from 0K to 310K in NVT ensemble with Nosé-Hoover thermostat^58^. Then we performed a single long equilibration for lipid and solvent (1000 ns) in NPT ensemble. During the heating step and the long equilibration, position restraints were placed of 10 kcal/mol-Å^2^ applied on the receptor, propionate and POPC/CHSD for the first 1 ns. Later, the restraint on lipids was reduced from 5 kcal/mol-Å^2^ to 0 kcal/mol-Å^2^ in steps of 1 kcal with 5 ns of simulations per step. Then the POPC/CHSD were allowed to freely move during the rest of the long equilibration and the final snapshot was used as the initial conformation for equilibrating the protein and ligand. The position restraints were applied on the protein (backbone and side chain) and ligand starting at 5 kcal/mol-Å^2^ reducing to 0 kcal/mol-Å^2^ in steps of 1 kcal/mol-Å^2^ with 5 ns of simulation per step. The last snapshot of the equilibration step was used as initial conformation for five production runs with random seeds. This snapshot was also used as reference conformation for all the RMSD in coordinates. The pressure was controlled using Parrinello-Rahman method^59^ and the simulation system was coupled to 1 bar pressure bath. In all simulations LINCS algorithm is applied on all bonds and angles of waters with 2 fs time step used for integration. We used a cut-off of 12 Å for non-bond interaction and particle mesh Ewald method^60^ to treat long range L-J interaction. The MD snapshots were stored at every 20 ps interval. Trajectories were visualized with VMD and PyMOL (Molecular Graphics System, Version 2.5 Schrödinger) and analyzed using the GROMACS package (version 2016/2019). All MD analysis was done on the aggregated trajectories from the 5 runs (total 5 × 1 μs = 5 μs). Heatmaps and other MD related plots were generated with Graphpad Prism 9, whereas structural figures were generated using PyMOL.

### Molecular dynamics analysis

#### Ligand-receptor and intramolecular interactions

Contact frequencies were calculated using the “get_contacts” module (https://getcontacts.github.io/). The following interaction types were calculated between ligand and receptor: hydrogen bonds, hydrophobic and van der Waals interactions.

#### Calculation of Residue Distances

For the distance between two residues, we used *gmx mindist* (GROMACS package 2016/2019), which calculates the minimal distance between two atoms (e.g., sidechain, Cα, oxygens, nitrogens) of one of each residue over time. Distance analysis on the static structures were done using the measurement tool in PyMOL. Chosen atoms for distance calculations are described in each legend.

#### Rotamer Analysis of F250

For the rotamer analysis of residues of interest, we used the VMD tcl script “Calculate_dihedrals” (https://github.com/aiasja/calculate_dihedrals).

#### Conformational Clustering

To select representative snapshots from MD simulations that are shown in Fig. 4i and Fig. 5b, we clustered (*gmx cluster*, GROMACS package 2016/2019) the aggregated trajectories using transmembrane helix backbone atoms. An RMSD cutoff for clustering was set at 0.08 nm for propionate-bound simulations, 0.085 nm for no-ligand WT simulations and 0.085 nm for no-ligand Q181D simulations. For propionate clustering (Fig. 2f), we used an RMSD cut-off of 0.01 nm for the ligand.

#### Root-Mean-Square Deviation (RMSD)

The *gmx rms* (GROMACS package 2016/2019) function was used to determine whether simulations were stableFor this we used the transmembrane backbone of OR51E2 by selecting the following residues: 23-50 (TM1), 57-86 (TM2), 93-126 (TM3), 137-164 (TM4), 191-226 (TM5), 230-264 (TM6), and 269-294 (TM7). As reference, we used the equilibrated MD structure of propionate bound, apo and Q181D OR51E2. In order to assess the stability of the ligand in the binding pocket over time, the RMSD of propionate was calculated using the equilibrated MD structure of propionate-bound as a reference.

#### Volume of the ligand binding pocket

The volume and surface area of the propionate binding pocket in OR51E2 was calculated using the Maestro SiteMap module^61,62^. Three structures were used for the volume calculation: 1) the OR51E2 cryo-EM structure bound to propionate, 2) the OR51E2-L158A model bound to hexanoate, 3) the OR51E2-F155A model bound to octanoate. To prepare the L158A and F155A models we used the Maestro mutation function to introduce the substitutions onto the cryo-EM structure of OR51E2; these models were then energy minimized using the ProteinPreparationWizard module using default parameters^56^. We then used Maestro Glide Docking^63–65^ to dock hexanoate and octanoate into the resulting models of OR51E2-L158A and OR51E2-F155A, respectively. We prepared the docking grid box for both OR51E2-L158A and OR51E2-F155A by defining a box centered at propionate, with a box length of 2.5 nm. Glide ligand docking was performed using XP precision and default parameters to yield a model for OR51E2-L158A bound to hexanoate and OR51E2-F155A bound to octanoate. To calculate ligand binding site volumes using the SiteMap module, we defined the ligand binding pocket as a maximum of 0.6 nm around selected ligand (propionate/hexanoate/octanoate) with at least 15 site points (probes) per reported site. The grid size for the probes was set to 0.035 nm. Using this approach, the calculated volumes for wild-type OR51E2, OR51E2-L158A, and OR51E2-F155A were 31 Å^3^, 68 Å^3^, and 90 Å^3^, respectively.

### Phylogenetic tree

A phylogenetic tree of human Class A GPCRs was made by analyzing 677 sequences. Of these, 390 sequences were from olfactory receptors (56 Class I ORs and 334 Class II ORs), while 287 were from non-olfactory Class A GPCRs. Sequences were aligned with Clustal^66^ on Jalview 2.11.2.5^67^. On R studio 202.07.01, alignment reading and matrix of distance between sequences (by sequence identity) calculation were performed with the BiostrinG_s_^68^ and seqinr^69^ packages. Neighbor-Joining tree and tree visualization were realized with packages ape^70^ and ggtree^71^ and the tree is plotted unrooted with the daylight method.

## Data Availability

Coordinates for propionate OR51E2-G_s_ have been deposited in the RCSB PDB under accession code 8F76. EM density maps for OR51E2-G_s_ and the 7TM domain of OR51E2 have been deposited in the Electron Microscopy Data Bank under accession codes EMD-28896, and EMD-28900, respectively. The MD simulation trajectories have been deposited in the GPCRMD database.

## Acknowledgements

We thank Dan Toso at Cal-Cryo at QB3-Berkeley for help in microscope operation and data collection. H.M., C.A.D.M. and J.T. thank Mengjue J. Ni and Hsiu-Yi Lu for their technical support. This work was supported by the National Institutes of Health (NIH) grant R01DC020353 (H.M., N.V., and A.M.) and K99DC018333 (C.A.D.M.). Cryo-EM equipment at UCSF is partially supported by NIH grants S10OD020054 and S10OD021741. This project was funded by the UCSF Program for Breakthrough Biomedical Research, funded in part by the Sandler Foundation. A.M. acknowledges support from the Edward Mallinckrodt, Jr. Foundation and the Vallee Foundation.

## Contributions

C.B.B., C.A.D.M., W.J.C.v.d.V., N.V., H.M., and A.M. designed the study. C.B.B. cloned constructs, prepared baculoviruses, expressed and purified G protein complexing reagents, and optimized large scale production of OR51E2. C.B.B. worked out conditions to biochemically purify and stabilize the propionate-bound OR51E2-G_s_ complex, and identified optimal cryo-EM grid preparation procedures following screening, collection, and processing of 200 kV cryo-EM data. B.F. and A.M. performed 300 kV cryo-EM data collection. C.B.B. determined high-resolution cryo-EM maps by extensive image processing with input from A.M. A.M. and C.B.B. built, refined models of propionate-bound OR51E2 in complex with G_s_ and Nb35. C.B.B. and A.M. analyzed cryo-EM data and models and prepared figures and tables. C.A.D.M. and J.T. analyzed OR models and sequences to design and clone OR mutants, performed Glosensor signaling experiments for OR functional activity, and generated OR cell surface expression data by flow cytometry with input from H.M. C.A.D.M and J.T. analyzed and prepared figures and tables for signaling and flow cytometry data. C.A.D.M. built the phylogenetic tree of ORs and non-olfactory Class A GPCRs. N.M. set up molecular dynamics simulations, ligand docking, and performed binding pocket volume calculations. W.J.C.v.d.V. analyzed simulation trajectories and prepared figures describing simulation data. W.J.C.v.d.V., N.M. and N.V. provided mechanistic insight from simulation data. C.L.D.T. performed bioinformatic analysis of OR and non-olfactory Class A GPCR conservation. L.L. and C.B.B. performed pilot GloSensor signaling studies in suspension cells. C.B.B., C.A.D.M., and A.M. wrote an initial draft of the manuscript and generated figures with contributions from all authors. Further edits to the manuscript were provided by W.J.C.v.d.V., N.M., V.N., and H.M. The overall project was supervised and funded by N.V., H.M., and A.M.

