## Supplementary Information for "Structural basis of odorant recognition by a human odorant receptor"

**Supplementary Figures/Tables:**

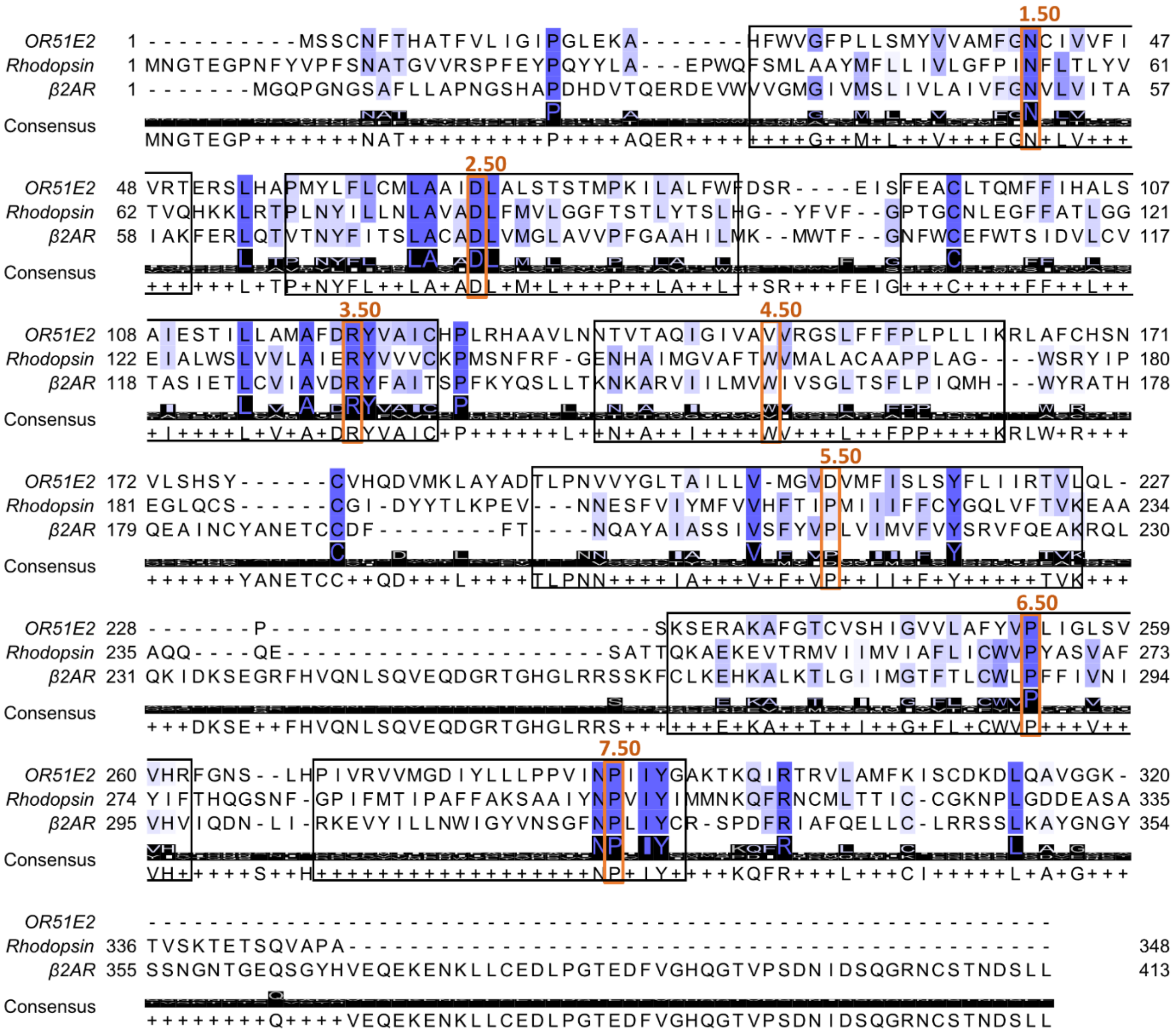


**Supplementary Fig. 1**. Alignment of OR51E2, rhodopsin and β2 adrenergic receptor (β2AR) amino acid sequences. Conservation is highlighted from low (white) to high (dark blue) and the consensus amino acid is shown. Transmembrane domains are boxed and labeled in black. The most conserved residue in class A GPCRs for each transmembrane domain is boxed and labeled in orange along with the Ballesteros-Weinstein number.

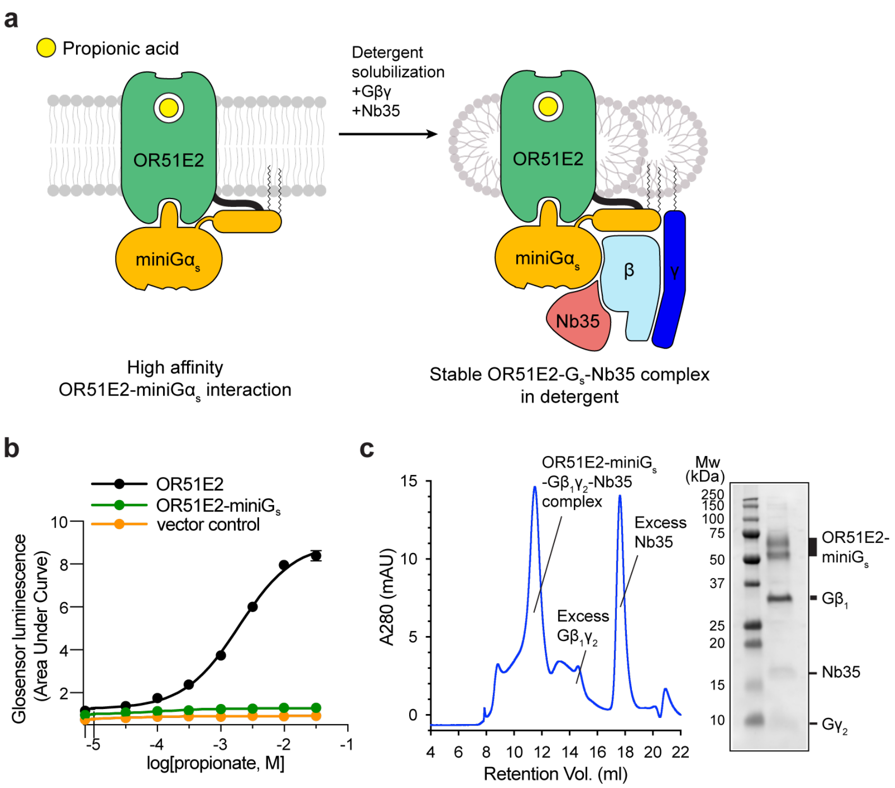


**Supplementary Fig. 2. Biochemical preparation of OR51E2-G$^s$ complex bound to propionate. a)** Schematic outlining the strategy for stabilization and purification of the activated OR51E2-G$_s$ complex bound to propionate**. b)** GloSensor cAMP assay demonstrating that fusion of miniG$_s$ to OR51E2 blocks activation of endogenous G$_s$ in response to treatment with propionate, suggesting that miniG$_s$ couples to the OR51E2 transmembrane core. Data points are analytical replicates from a representative experiment. **c)** Size-exclusion chromatogram of purified OR51E2-G$_s$-Nb35 complex used for structure determinations shown together with a representative SDS-PAGE gel analysis of the collected fraction containing the OR51E2-G$_s$-Nb35 complex.

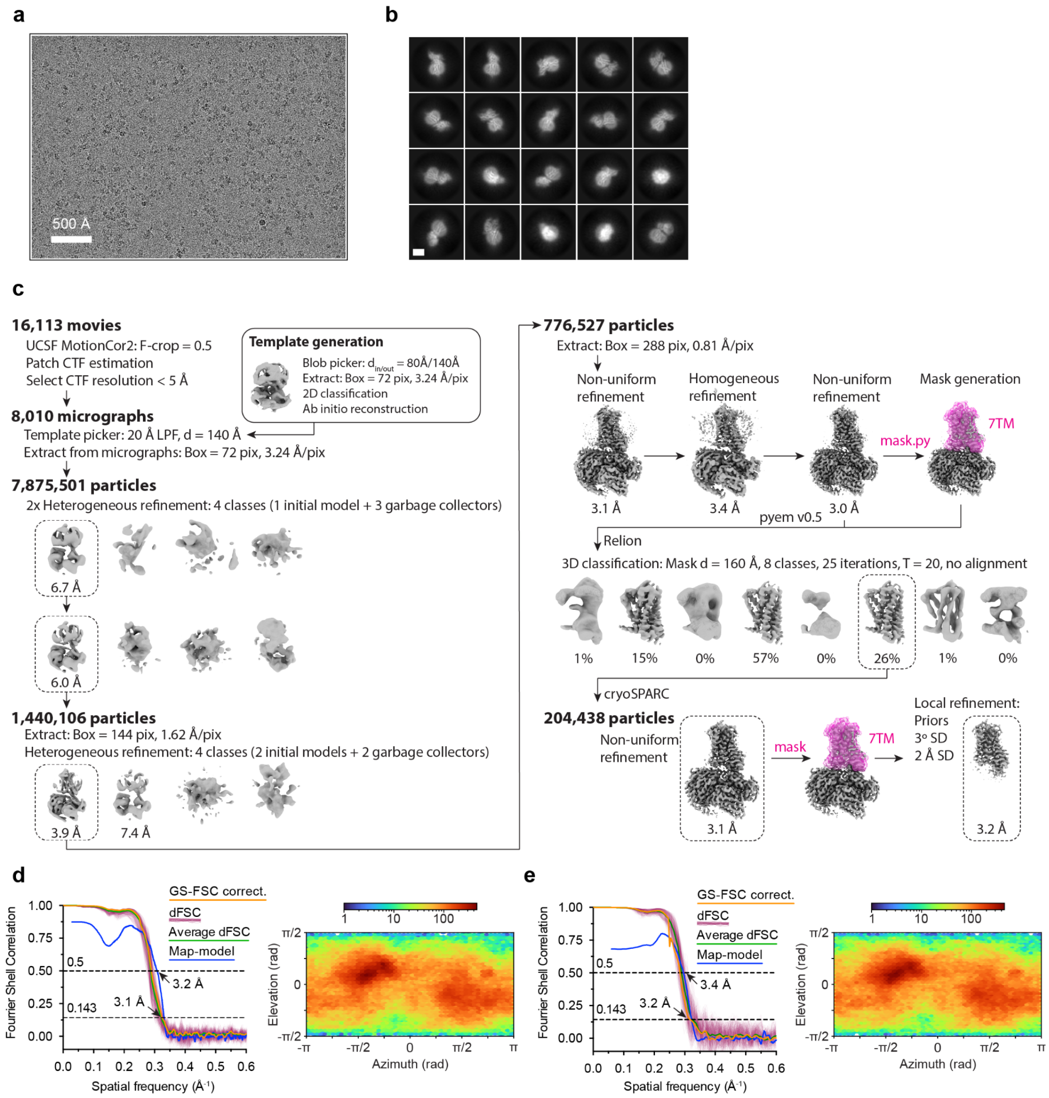


**Supplementary Fig. 3. Cryo-EM data processing for OR51E2-G$_s$. a)** A representative cryo-EM micrograph from the curated OR51E2-G$_s$ dataset (n = 8,010) obtained from a Titan Krios microscope. **b)** A subset of highly populated, reference-free 2D-class averages are shown. Scale bar is 50 Å. **c)** Schematic showing the image processing workflow for OR51E1-G$_s$. Initial processing was performed using UCSF MotionCor2 and cryoSPARC. Particles were then transferred using the pyem script package[39] to RELION for alignment-free 3D classification. Finally, particles were processed in cryoSPARC using the non-uniform and local refinement tools. Dashed boxes indicate selected classes, and 3D volumes of classes and refinements are shown along with global Gold-standard Fourier Shell Coefficient (GSFSC) resolutions. **d, e)** Map validation for the OR51E2-G$_s$ (**d**) globally refined, and (**e**) locally refined cryo-EM maps. Gold-standard Fourier shell correlation (FSC) curves are calculated in cryoSPARC, and shown together with directional FSC (dFSC) curves generated with dfsc.0.0.1.py as previously described[40]. Map-model correlations calculated in the Phenix suite are also shown. Arrows indicate map and map-model resolution estimates at 0.143 and 0.5 correlation respectively. Euler angle distributions calculated in cryoSPARC are also provided for each map.

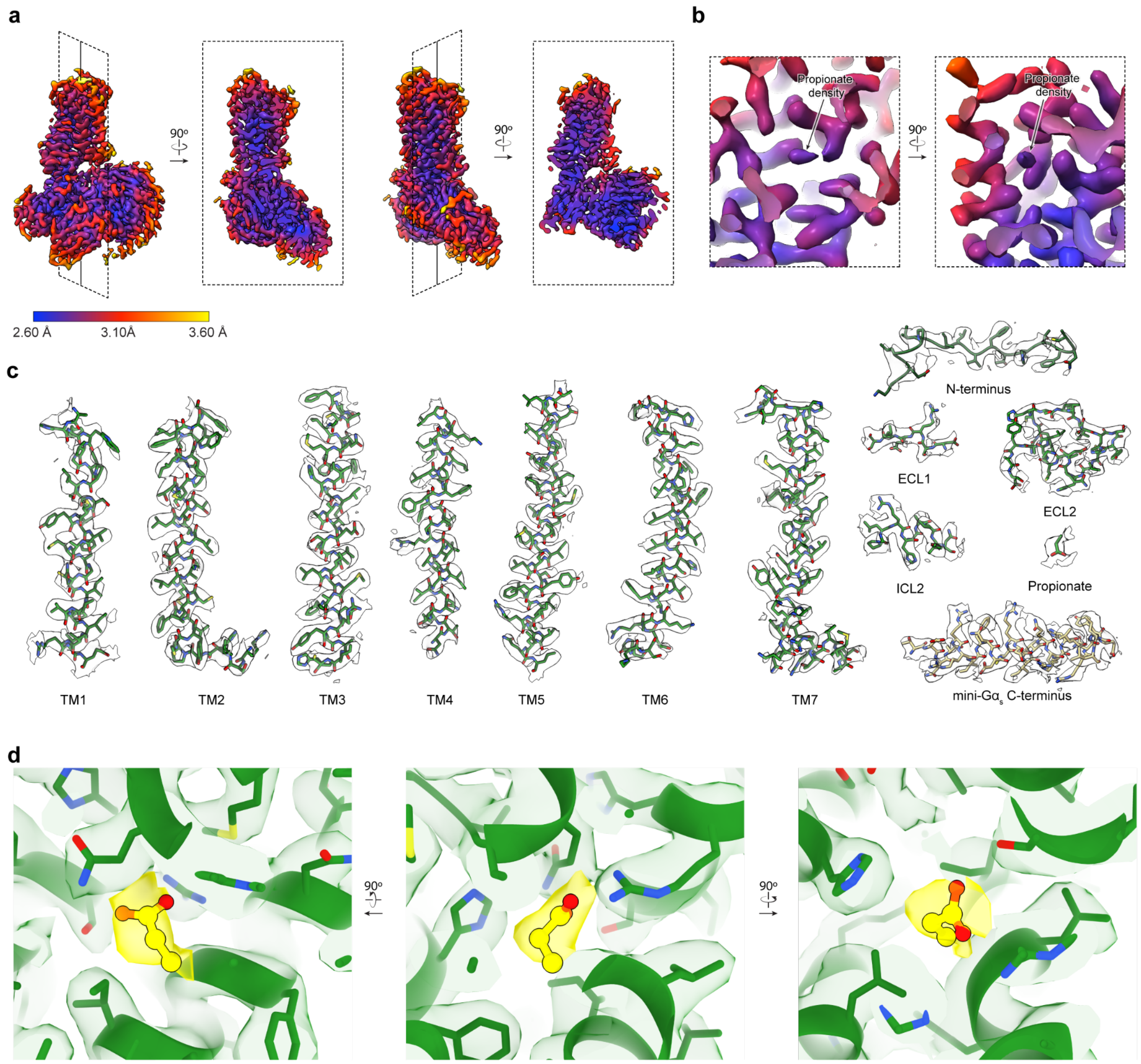


**Supplementary Fig. 4. Cryo-EM density and atomic model. a)** Orthogonal views of local resolution map of OR51E2-$G_s$ calculated with the local resolution estimation tool in cryoSPARC. **b)** Close-up view showing the local resolution of the propionate binding site. **c)** Representative cryo-EM densities from the 3D reconstruction of OR51E2. Shown are the transmembrane helices and loop regions of OR51E2 as well as the C-terminal helix of miniG$\alpha_s$. **d)** Close-up view of cryo-EM density (yellow sticks and density) supporting propionate binding pose.

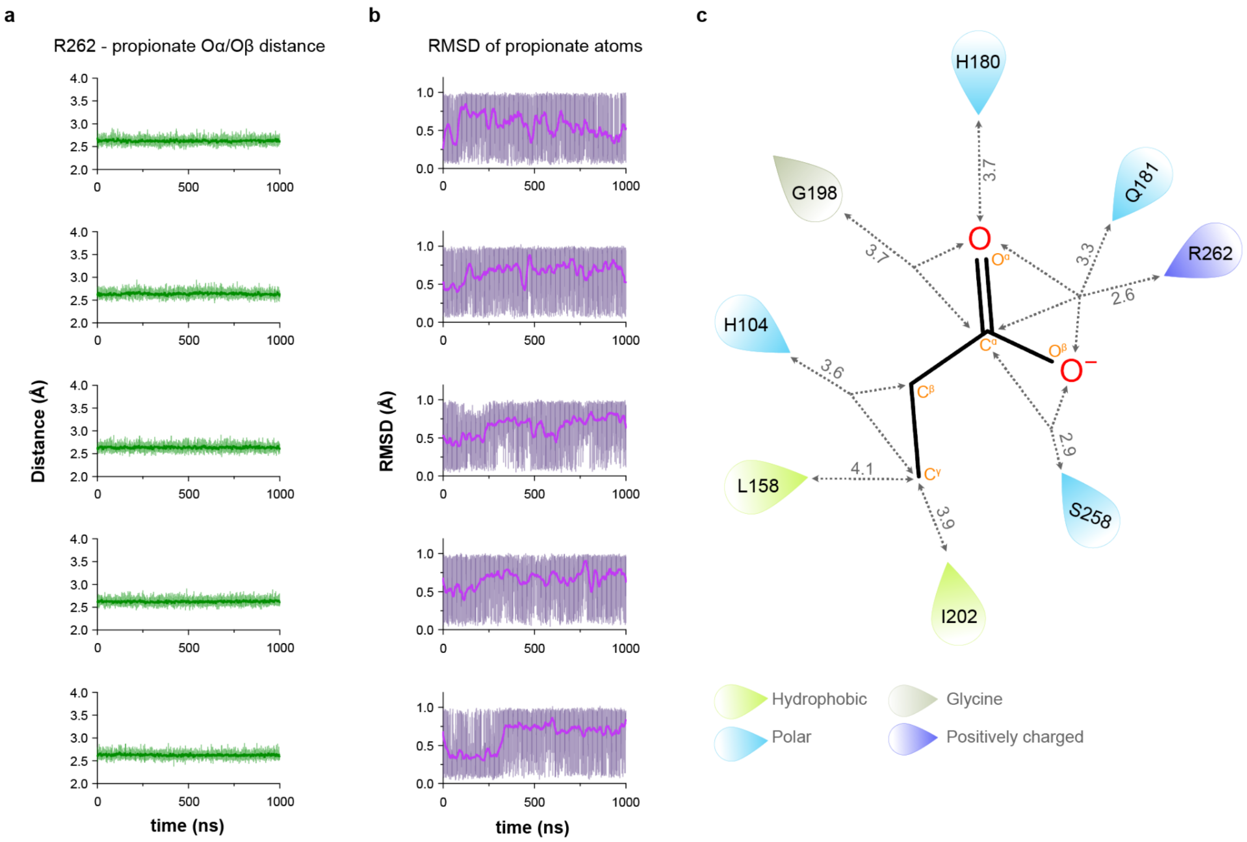


**Supplementary Fig. 5. Interactions between propionate and OR51E2 in molecular dynamics simulations. a)** Distance plot between R262 and propionate. Distance was measured between nitrogens of R262 and oxygens of propionate. **b)** Root-mean-square deviation **(**RMSD) values of production simulation runs for propionate calculated with reference to the equilibrated structure of OR51E2 prior to 1 μs production simulation. **c)** Minimal distances between ligand atoms and residues are shown in gray (Å). Gray dashed arrows highlight which interactions are made between a certain receptor residue and ligand atom(s) (see figure 2G). All data are shown as means from n = 5 independent runs (at different velocities) each 1 μs long. Standard deviation of measurement for each of the residue-ligand distance are as follows; 0.03 Å (R262), 0.10 Å(S258), 0.16 Å (I202), 0.12 Å (G198), 0.23 Å (Q181), 0.23 Å (H180), 0.25 Å (L158), and 0.14 Å (H104).

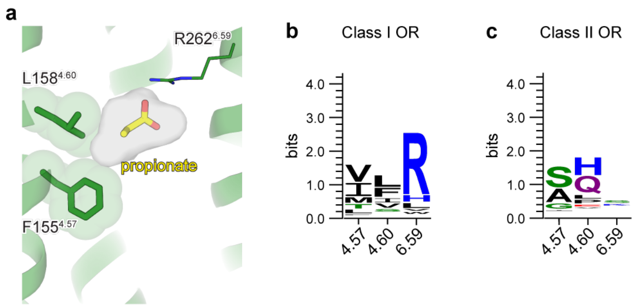


**d**

| | Class I OR | | | | Class II OR | | |
|---|---|---|---|---|---|---|---|
| | 4.57 | 4.60 | 6.59 | | 4.57 | 4.60 | 6.59 |
| A | 0 | 0 | 2 | A | 28.7 | 1.2 | 8.4 |
| C | 0 | 4 | 2 | C | 0 | 0.3 | 0.9 |
| D | 0 | 2 | 0 | D | 0.3 | 1.8 | 1.5 |
| E | 2 | 0 | 0 | E | 0.3 | 4.8 | 1.2 |
| F | 4 | 23 | 2 | F | 1.5 | 0.9 | 7.8 |
| G | 0 | 0 | 0 | G | 13.1 | 0.3 | 5.7 |
| H | 2 | 0 | 9 | H | 0 | 31 | 4.2 |
| I | 27 | 13 | 0 | I | 4.2 | 0.9 | 0.9 |
| K | 0 | 0 | 0 | K | 0 | 1.8 | 14.6 |
| L | 7 | 29 | 5 | L | 0.6 | 8.1 | 4.5 |
| M | 13 | 2 | 0 | M | 0.3 | 0.9 | 0.3 |
| N | 0 | 0 | 0 | N | 0 | 1.5 | 4.8 |
| P | 0 | 4 | 0 | P | 2.7 | 7.8 | 8.1 |
| Q | 0 | 0 | 2 | Q | 0.3 | 30.7 | 3 |
| R | 0 | 0 | 71 | R | 0 | 1.8 | 6.6 |
| S | 2 | 9 | 0 | S | 43 | 0.3 | 21.2 |
| T | 11 | 2 | 0 | T | 2.1 | 0.9 | 4.5 |
| V | 34 | 13 | 4 | V | 2.4 | 3.3 | 2.1 |
| Y | 0 | 2 | 0 | Y | 0.6 | 1.8 | 0 |
| W | 0 | 0 | 4 | W | 0 | 0 | 0 |

**Supplementary Fig. 6. Conservation of residues within the odorant binding pocket. a**) View of propionate-contacting residues contacting. Conservation weblogo of key residues in Class I (**b**) and Class II ORs (**c**). **d**) Specific amino acid frequencies at each position are shown for Class I and Class II ORs. OR51E2 residues at each position are indicated by a black box.

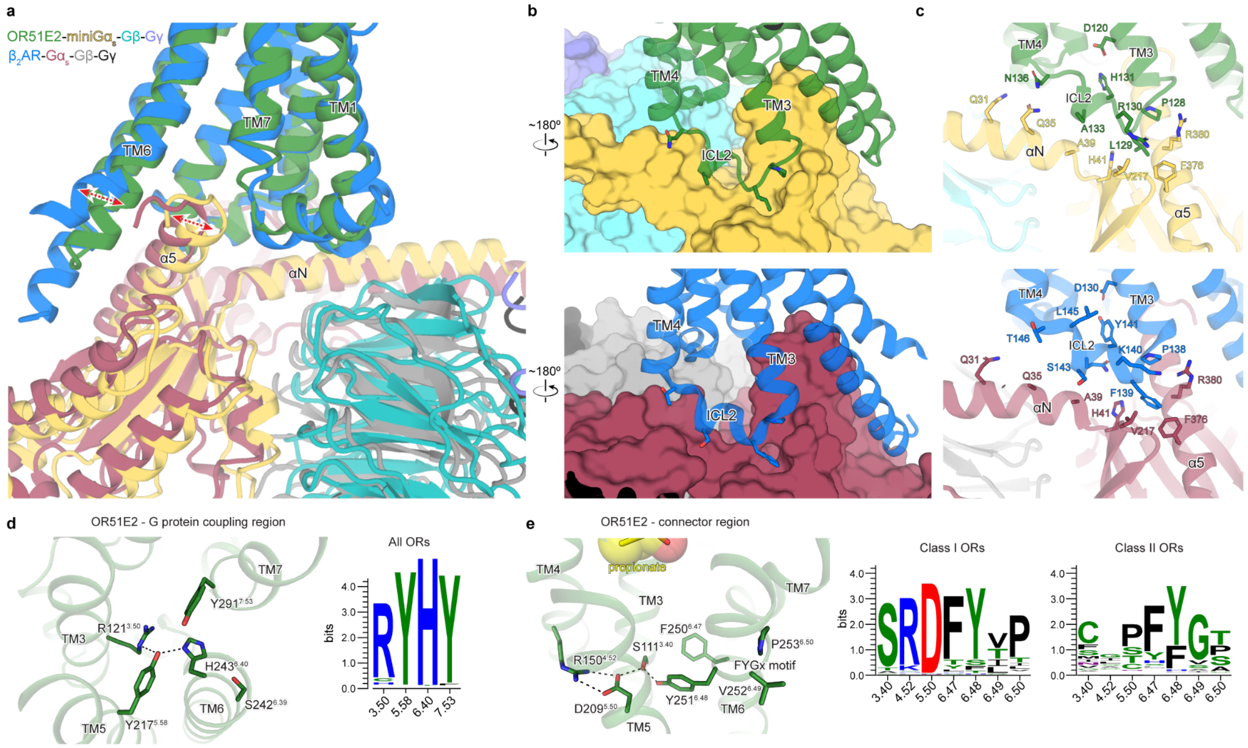

**Supplementary Fig. 7. Analysis of active state structure of OR51E2. a)** Structural comparison of G protein interaction for OR51E2 (green) and β2-adrenergic receptor (β2AR in blue, PDB code: 3SN6). **b)** Close-up views of intracellular loop 2 (ICL2) interaction with the $G\alpha_s$ subunit shown in surface representation. **c)** interactions between residues in ICL2 and the αN and α5 helices of the $G\alpha_s$ subunit. **d)** G protein-coupling region of OR51E2 is shown along with a weblogo (right) highlighting conservation of key residues for all human ORs. **e)** Residues that participate in the extended interaction hydrogen bonding network between TM3, TM4, TM5, and TM6 are conserved in human Class I ORs, but not in Class II ORs.

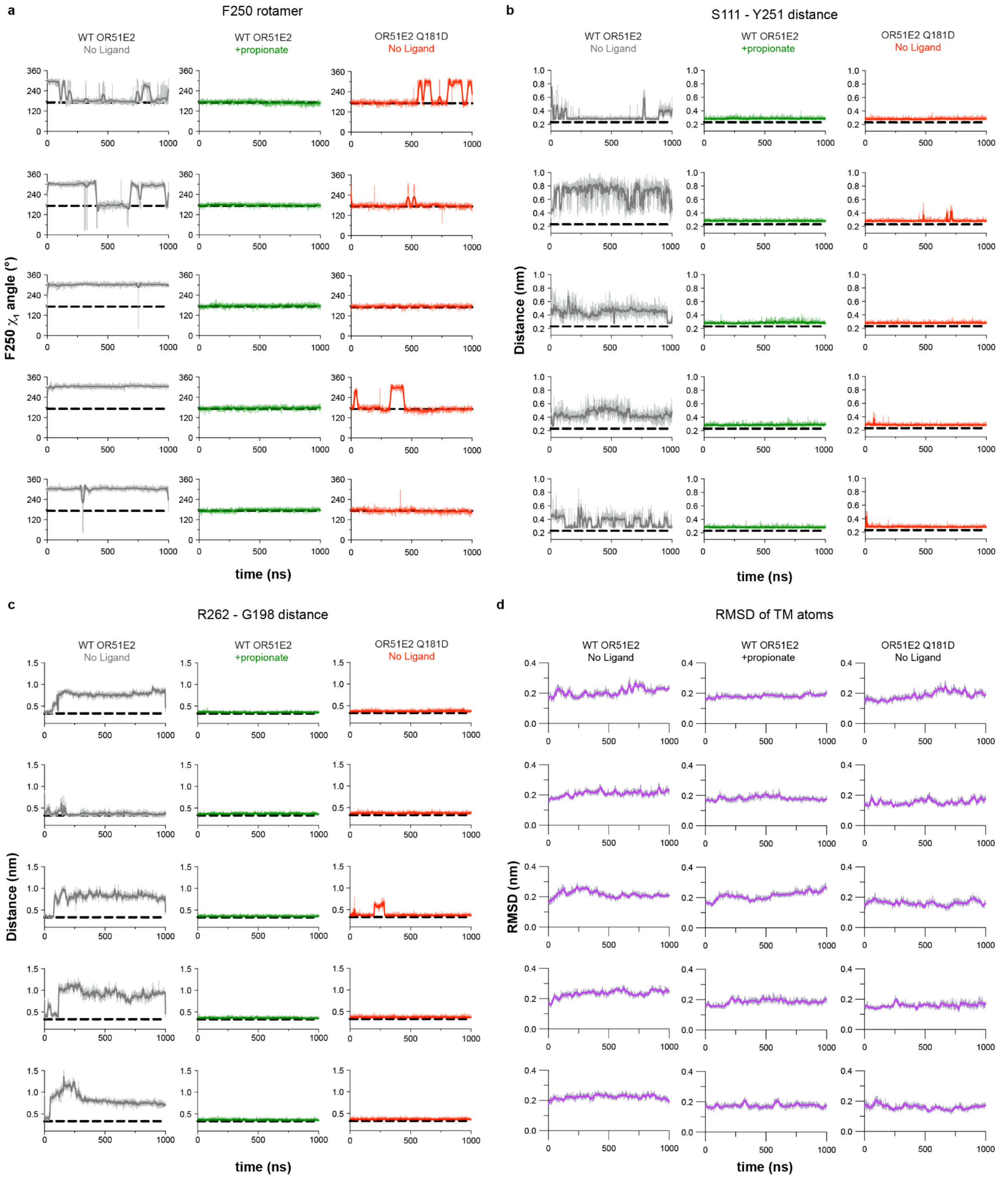


**Supplementary Fig. 8. MD simulation trajectory results. a)** $\chi$1 angle of F250 for WT and Q182D OR51E2. **b)** Distance plot between S111 and Y251 with simulation time. Distance was measured between oxygen atoms of the hydroxyl groups in the side chains of S111 and Y251. **c)** Distance plot between R262 and G198 as a function of simulation time for WT and Q181D OR51E2. The minimum distance was measured between R262 sidechain atoms and G198 mainchain atoms (excluding the hydrogens). **d)** Root-mean-square deviation (RMSD) values for TM backbone atoms in the transmembrane helices (*see methods*) calculated with reference to the equilibrated structure of the apo and propionate bound OR51E2 simulations, as well as for simulations of Q181D OR51E2. All data are shown as five independent runs (at different velocities) each 1 μs long for no-ligand (gray), propionate bound (green) and Q181D no-ligand (red).

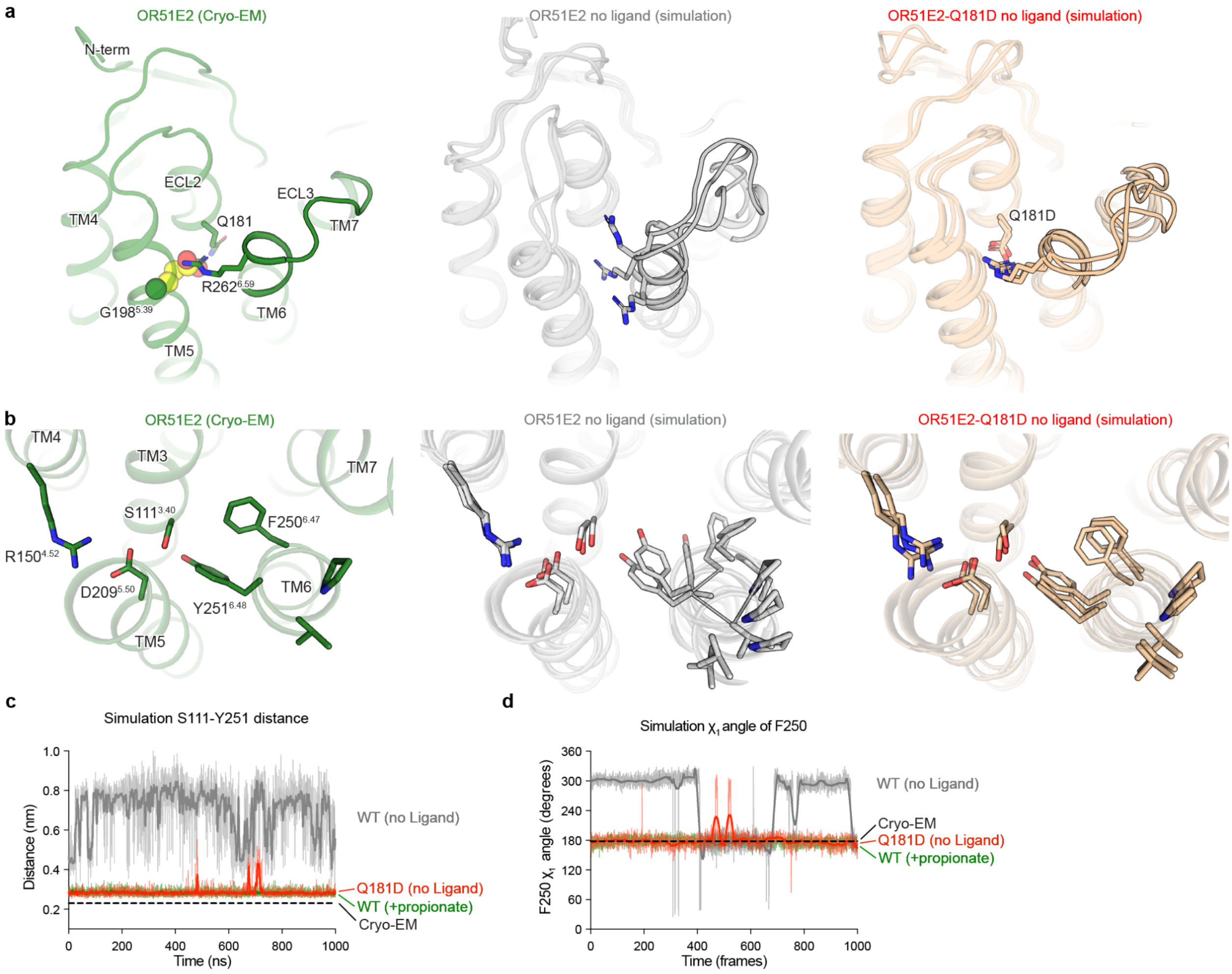


**Supplementary Fig. 9. Hallmarks of ligand-independent activation in Q181D.**

**a and b**) Comparison of cryo-EM structure of propionate-bound OR51E2 with snapshots from molecular dynamics (MD) simulations of WT OR51E2 with propionate removed, and OR51E2 Q181D. **a)** Close-up views of OR51E2 binding site and ECL3 region, shows that introduction of Asp in position 181 (Q181D) stabilizes R262$^{6.59}$ in an active-like state by a direct ionic interaction. **b**) Close-up views of OR51E2 connector region, suggests increased stabilization of TM6 for the Q181D mutant. **c and d**) Molecular dynamics trajectories from representative simulations to highlight structural organization of connector region. **c)** Distance between S111$^{3.40}$ and Y251$^{6.47}$ hydroxyl groups is comparable for Q181D and propionate-bound WT OR51E2. **d)** Rotamer angle of F250$^{6.47}$ is comparable for Q181D and propionate-bound WT OR51E2. Simulations were performed with or without propionate over the course of 1000 ns (see Supplementary Fig. 8 for replicates of simulation trajectories).

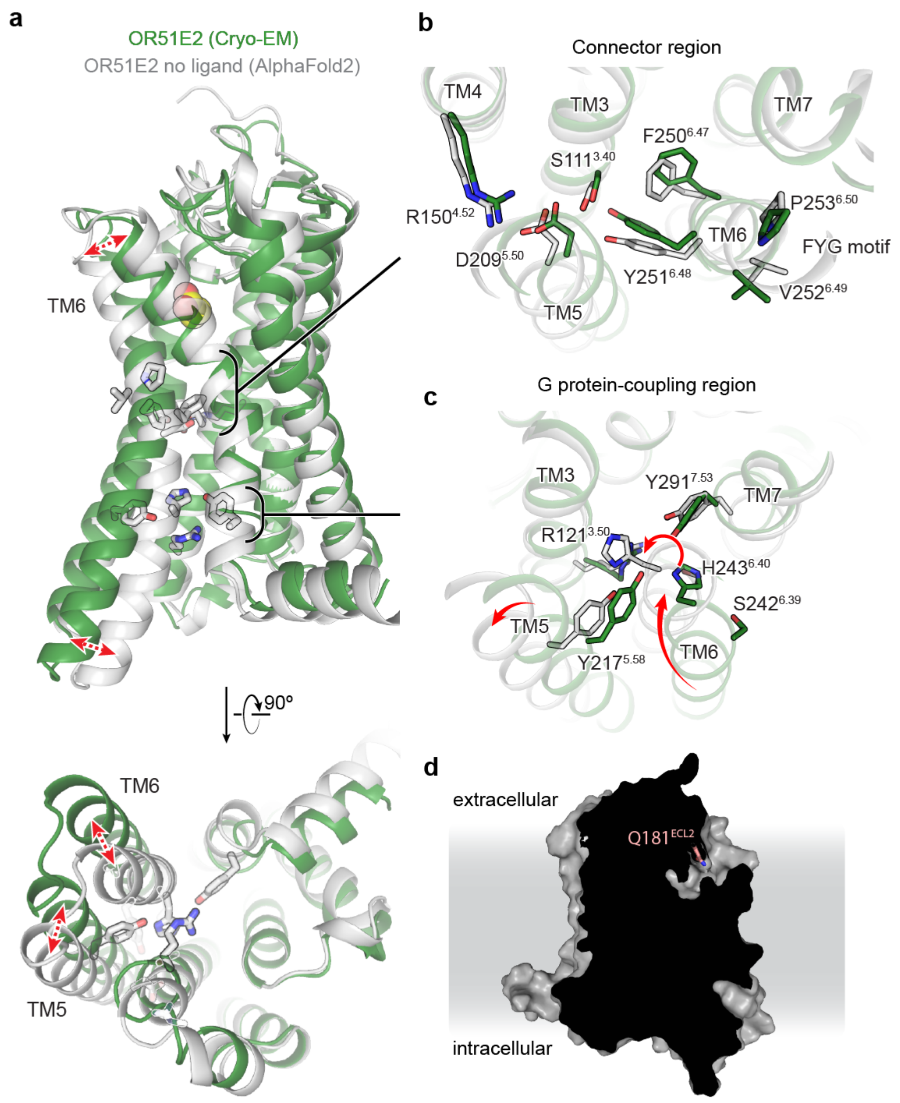


**Supplementary Fig. 10. AlphaFold2 model of OR51E2. a)** AlphaFold2 predicted structure of unbound OR51E2 (gray) superimposed onto the experimentally determined structure of propionate-bound OR51E2 in the active state (green cartoon and yellow spheres). In the AlphaFold2 model, TM6 is inwardly displaced compared to the active structure. Closeup views of (**b**) the Connector region and (**c**) the G protein-coupling region are provided. **d**) Slice through surface representation of AlphaFold2 predicted OR51E2, suggests solvent accessibility of the ligand binding site in the inactive state.

**Extended Data Table 1. Cryo-EM data collection, refinement, and validation statistics**

| | **Propionate-bound OR51E2-$G_s$** |
|---|---|
| EMDB: Full map | EMD-28896 |
| EMDB: 7TM map | EMD-28900 |
| RCSB PDB: Model | 8F76 |
| **Data collection** | |
| Microscope | Thermo Scientific Krios G3i |
| Detector | Gatan K3 with Gatan BioQuantum Energy filter |
| Voltage (kV) | 300 |
| Magnification | 105,000 |
| Defocus range (μm) | -1.0 to -2.1 |
| Pixel size, physical (Å) | 0.81 |
| Total exposure ($e^-$/Å$^2$) | 50 |
| Frame exposure ($e^-$/Å$^2$/frame) | 0.833 |
| Images, number of | 16,113 |
| Frames/image, number of | 60 |
| Initial particles, number of | 7,875,501 |
| Final particles, number of | 204,438 |
| Symmetry imposed | C1 |
| Map sharpening, *B* factor (Å$^2$) | |
| Full map | -140.2 |
| 7TM map | -162.8 |
| Map resolution, masked (Å) | |
| Full map | 3.1 |
| 7TM map | 3.2 |
| FSC threshold | 0.143 |
| **Refinement** | |
| Initial model used (AlphaFold code) | Q9H255 |
| Model resolution (Å) | 3.2 |
| FSC threshold | 0.5 |
| Model composition | |
| Chains | 6 |
| Non-hydrogen atoms | 8,176 |
| Protein residues | 1,038 |
| Ligands | 1 |
| *B* factors (Å$^2$) | |
| Protein | 36.53 |
| Ligand | 37.12 |
| R.m.s. deviations | |
| Bond length (Å) | 0.004 |
| Bond angles (°) | 1.022 |
| Validation | |
| MolProbity score | 1.44 |
| Clash score | 4.84 |
| EMRinger score | 3.66 |
| Rotamer outliers (%) | 0.23 |
| Ramachandran plot | |
| Favored (%) | 96.86 |
| Allowed (%) | 3.14 |
| Disallowed (%) | 0.00 |

**Extended Data Table 2. Expression and pharmacodynamic constants for OR51E2 variants.**

| | Surface expression (mean) | *Propionate activity in GloSensor cAMP assay* | | | |
|---|---|---|---|---|---|
| | | $EC_{50}$ (mM, mean) | $pEC_{50}$ (-log M, mean ± S.E.M.) | Emax (mean ± S.E.M.) | Activity index (Emax * $pEC_{50}$) |
| **WT** | 8,490 | 0.824 | 3.08 ± 0.053 | 3.24 ± 0.049 | 9.98 |
| **H104A** | 4,130 | 17.1 | 1.77 ± 0.77 | 1.74 ± 0.63 | 3.08 |
| **S111A** | 112 | n.r. | n.r. | n.r. | n.r. |
| **R121A** | 192 | n.r. | n.r. | n.r. | n.r. |
| **R150A** | 163 | n.r. | n.r. | n.r. | n.r. |
| **F155A** | 8,700 | 12.3 | 1.91 ± 0.77 | 2.51 ± 0.099 | 4.79 |
| **L158A** | 5,250 | 1.6 | 2.8 ± 0.42 | 3.05 ± 0.43 | 8.53 |
| **H180A** | 4,570 | n.r. | n.r. | n.r. | n.r. |
| **Q181A** | 7,340 | 12.4 | 1.91 ± 0.32 | 2.68 ± 0.49 | 5.11 |
| **Q181D** | 9,140 | 38.9 | 1.41 ± 0.12 | 2.59 ± 0.25 | 3.65 |
| **Q181E** | 8,750 | n.r. | n.r. | n.r. | n.r. |
| **Q181N** | 4,910 | 3.9 | 2.41 ± 0.11 | 1.15 ± 0.016 | 2.77 |
| **G198A** | 5,960 | n.r. | n.r. | n.r. | n.r. |
| **I202A** | 9,730 | 13.7 | 1.86 ± 0.050 | 2.25 ± 0.060 | 4.2 |
| **D209A** | 112 | n.r. | n.r. | n.r. | n.r. |
| **Y217A** | 97.6 | n.r. | n.r. | n.r. | n.r. |
| **S242A** | 10,300 | 0.615 | 3.21 ± 0.080 | 3.48 ± 0.071 | 11.2 |
| **H243A** | 260 | 0.459 | 3.34 ± 0.095 | 1.39 ± 0.015 | 4.64 |
| **F250H** | 2,420 | 4.12 | 2.39 ± 0.036 | 3.01 ± 0.066 | 10.2 |
| **F250Y** | 3,450 | 0.423 | 3.37 ± 0.096 | 2.68 ± 0.039 | 6.38 |
| **Y251A** | 114 | n.r. | n.r. | n.r. | n.r. |
| **Y251F** | 1,500 | 4.72 | 2.33 ± 0.031 | 2.39 ± 0.029 | 5.56 |
| **Y251H** | 254 | n.r. | n.r. | n.r. | n.r. |
| **V252A** | 6,610 | 1.01 | 3.00 ± 0.085 | 2.97 ± 0.076 | 8.91 |
| **P253A** | 1,460 | 5.53 | 2.26 ± 0.05 | 2.32 ± 0.047 | 5.23 |
| **S258A** | 7,230 | n.r. | n.r. | n.r. | n.r. |
| **R262A** | 9,290 | n.r. | n.r. | n.r. | n.r. |
| **Y291A** | 3,080 | 1.4 | 2.85 ± 0.042 | 2.50 ± 0.031 | 7.13 |
| **mock** | 49.4 | n.r. | n.r. | n.r. | n.r. |

*n.r. = no response (fit $R^2 < 0.90$)*
